# mRNA decay can be uncoupled from deadenylation during stress response

**DOI:** 10.1101/2023.01.20.524924

**Authors:** Agnieszka Czarnocka-Cieciura, Matti Turtola, Rafał Tomecki, Paweł Krawczyk, Seweryn Mroczek, Wiktora Orzeł, Upasana Saha, Torben Heick Jensen, Andrzej Dziembowski, Agnieszka Tudek

## Abstract

The polyadenosine tail (pA-tail) regulates mRNA nuclear export, stability, and translatability. Based on reporter constructs, the prevailing model suggests that pA-tail removal mediated by Ccr4-NOT or PAN2/3 deadenylases is required for mRNA decapping and degradation. Here, we use direct RNA sequencing to track mRNA deadenylation and decay at steady-state and in stress conditions to show a global correlation between deadenylation and decay. Interestingly, codon optimality, previously postulated to dictate mRNA stability, only strongly affects decay of conserved and abundant transcripts, such as coding for ribosomal protein subunits. Degradation of those mRNAs is also accelerated in response to stress. Still, the in-depth analysis revealed that deadenylation is a factor that contributes to degradation but is not indispensable for decapping. We further demonstrate that deadenylation is the fastest for newly made tails depending on polyA-binding protein Pab1. Unexpectedly, decapping initiates on mRNAs of pA-tails of 20-35 adenosines presumably bound by Pab1.

## INTRODUCTION

Synthesis of the 3’ end polyadenosine tail (pA-tail) is a crucial step in mRNA biogenesis which regulates both nuclear and cytoplasmic aspects of transcript metabolism in all eukaryotes. In budding yeast, the Pap1 polyA-polymerase produces *de novo* 60 adenosine-long pA-tails, in a process which is co-regulated by the nuclear polyA-binding protein Nab2 and the Cleavage and Polyadenylation Factor (CPF) complex (Turtola et al, 2021; Aibara et al., 2017; Rodriguez-Molina and Turtola 2022). At least 40 adenosine long pA-tails are required to both protect the transcript from nuclear decay and enable the mRNA to be exported to the cytoplasm (Dower et al., 2004). The Nab2 protein mediates the protective function of the pA-tail, in the absence of which nuclear transcripts are rapidly degraded (Schmid et al., 2015). mRNA export is mediated by the conserved Mex67-Mtr2 hetero-dimer (De Magistris, 2021). Inhibition of this process, achieved by depletion of Mex67 (Haruki et al., 2008), also leads to massive decay of newly made mRNAs due to sequestration of Nab2 on RNAs accumulated in the nucleus (Tudek et al., 2018).

Once the mRNA reaches the cytoplasm Nab2 and other nuclear factors are released, allowing Nab2 to be re-imported to the nucleus (Tran et al., 2007), and the transcript becomes associated with cytoplasmic proteins. Among those key factors are the polyA-binding protein Pab1 and the cap-binding eIF4E translation factor, which are thought to form a complex connecting both ends of the transcript. This looping of the mRNA was suggested to play a role in translation and decay regulation (Tarun et al., 1997; Otero et al., 1999; Archer et al., 2015).

During its lifetime in the cytoplasm, the mRNA pA-tail is gradually shortened in a process called deadenylation by the two main deadenylase complexes, PAN2/3 and Ccr4-NOT. In mammals, a bi-phasic deadenylation model has been proposed that postulates that decay of the pA-tail is initiated by PAN2/3 and taken over by Ccr4-NOT, with the caveat that PAN2/3 is actually nearly dispensable for deadenylation (Chen and Shyu, 2011; Yi et al., 2018). However, in budding yeast, each complex clearly has its preferred set of substrates (Tudek et al., 2021). Yeast PAN2/3 is recruited by binding to Pab1 multimers and the pA-tail (Schäfer et al., 2019; Wolf et al., 2014), and targets predominantly mRNAs of high abundance. Ccr4-NOT is recruited more efficiently to mRNAs of low abundance, most likely by the action of its sequence-specific Puf1-5 co-factors, though Ccr4-NOT was also suggested to bind Pab1 (Webster et al., 2018; Webster et al., 2019; Ulbricht and Olivas, 2008; Goldstrohm et al., 2006). Those discrepancies might stem from evolutionary changes in yeast and human cells, best pictured by the shift in the importance of Ccr4 and Pop2 Ccr4-NOT subunits for catalysis (Yi et al., 2018; Tucker et al., 2002; Webster et al., 2018).

*In vivo*, the mRNA pA-tails are coated by Pab1 polyA-binding protein, which is an important player in mRNA metabolism (Brambilla et al., 2019). *In vitro* data shows that Pab1 significantly contributes to pA-tail length regulation by stimulating deadenylation by Ccr4-NOT on a 60 A substrate and providing a temporary block on a 30 A RNA (Webster et al., 2018). Similar *in vitro* assays showed that PAN2/3, independently of Pab1, deadenylates long pA-tails more efficiently than the short ones of 20-25 As, and that the presence of Pab1 stimulates PAN2/3 activity, but is not required for deadenylation (Wolf et al., 2014). Despite the wealth of published *in vitro* assays, the role of Pab1 *in vivo* for the mRNA pA-tail metabolism has not been fully elucidated. Furthermore, the cellular coexistence of both PAN2/3 and Ccr4-NOT, for which Pab1 presence might have pleiotropic effects, prevents making general assumptions of the role of Pab1 in the regulation of the polyadenylated transcriptome metabolism based on *in vitro* assays, substantiating the need for a global *in vivo* approach.

The accepted models of mRNA decay, many of which were derived from reporter systems, state that substantial pA-tail shortening results in decapping by Dcp1/2 followed by Xrn1 5’-3’ exonuclease-mediated decay (De Magistris, 2021; Chen and Shyu, 2011; Yi et al., 2018; Decker and Parker, 1993). At precisely which pA-tail-length cap removal occurs remains unknown. It is also not clear how events at the 3’end pA-tail mechanistically facilitate decapping at the 5’end.

Here, we asked whether the widely accepted paradigm stating that deadenylation is a prerequisite to mRNA decay is supported by transcriptome-wide *in vivo* data. Using Nanopore Direct RNA Sequencing, we calculated deadenylation and decay rates for cells grown at permissive temperature and under stress conditions inflicted by thiolutin treatment or heat shock. We show that at steady-state growth conditions, deadenylation is transcriptome-wide dependent on Pab1 and occurs most rapidly when polyadenosine is newly-made. We also show that decay positively correlates with deadenylation rates at permissive growth conditions and that both processes are linked to codon optimality. Strikingly, even though deadenylation and decay are accelerated during stress response in a wild-type situation, in a double *ccr4Δ pan2Δ* mutant decay is not inhibited, but instead delayed. Thus, we propose that deadenylation acts as an auxiliary factor accelerating degradation under certain conditions. Accordingly, we show that decapping occurs close to steady-state mean pA-tail values of 20-30 adenosines and that some mRNAs are not deadenylated prior to decay or require a factor dependent on nuclear mRNA export for decapping. Altogether, this suggests that the signal for decapping often operates independently of any substantial reduction of pA-tail length.

## RESULTS

### mRNA nuclear export block reveals cytoplasmic deadenylation and decay dynamics

We decided to take advantage of the DRS method (Tudek et al., 2021) to investigate the relationship between mRNA deadenylation and decay rates for yeast mRNAs. Previous studies showed that bulk yeast mRNA half-lives are 12 minutes (Miller et al., 2011; Sun et al., 2012; Neymotin et al., 2014; Chan et al., 2018). Therefore, it was critical to efficiently decouple synthesis and decay within the shortest possible time frame. In our previous works (Tudek et al., 2018, Schmid et al., 2018), we showed that nuclear depletion of the main cellular export factor, Mex67, using the Anchor-Away system (Haruki et al., 2008) leads to massive degradation of all newly made mRNAs (Figure 1A) within the first minutes after export-block. Transcripts are likely degraded prior to adenylation by Pap1. There is only a small number of mRNAs, which escape decay, and they are mostly hyperadenylated (Turtola et al., 2021; Jensen et al., 2001), making it possible to distinguish them from species normally detected in wild-type cells. In contrast, already exported mRNAs continue their metabolism in the cytoplasm, allowing the monitoring of deadenylation and degradation kinetics for this pool of transcripts. To our knowledge, there is no other more efficient system to arrest mRNA production in such a short period of time. Using several Mex67-depletion time points we verified that this approach resulted in a marked decrease in global pA-tail lengths over time, which was mostly seen in the upper pA-tail length distributions (Figure 1B) and subsequently produced three replicates for Direct RNA Sequencing (Supplemental Figure 1A), which were all averaged to model decay and deadenylation transcriptome-wide.

**Figure 1.**
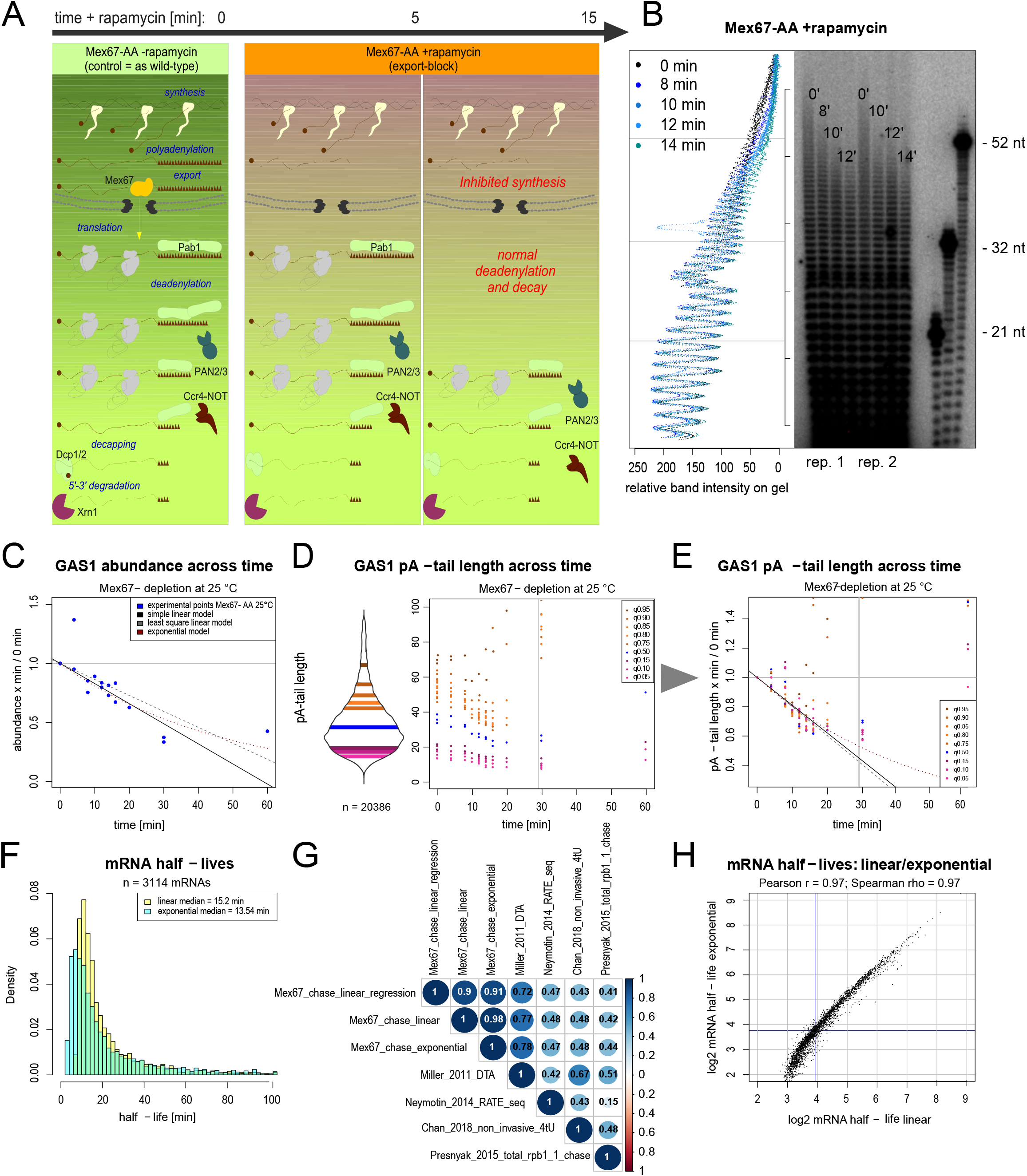
Overview of the experimental setup produced to model mRNA deadenylation and decay rates in yeast. **A**. Schematic representation of the phenotypes of Mex67-AA nuclear depletion and the utility of this genetic background to study cytoplasmic mRNA decay and degradation. **B**. Autoradiogram on the right side shows global distribution of pA-tail lengths in Mex67-depleted cells compared to control. The graph on the left side shows quantification of the autoradiogram lanes. **C**. Decay curve of *GAS1* mRNA produced from all three Mex67-AA replicates with transcript levels represented as a fraction of the control sample. The horizontal line at y = 1 shows the cutoff used to reject any y > 1 values for x > 0 min. Shown are three lines that describe decay of the transcript in a linear and exponential manner. More single gene examples are shown in Supplemental Figure 1. **D**. Left panel shows the distribution of pA-tail length in control cells for GAS1 mRNA. Right panel shows raw values of pA-tail length quantiles across three Mex67-AA 25 °C datasets. **E**. Graph showing a change in pA-tail quantile values across time in Mex67-AA 25 °C datasets by representing them as a fraction of the control quantile value. The horizontal line at y = 1 shows the cutoff used to reject any y > 1 values for x > 0 min, while the vertical line at x = 29 min shows that y values for x >29 min were not used for deadenylation rate calculation. Shown are three lines that describe decay of the transcript in a linear and exponential manner. **F**. Histogram showing distribution of mRNA half-lives calculated from the Mex67-AA 25 °C chase datasets using a linear and exponential model. **G**. Pairwise correlation matrix of mRNA half-lives obtained from the Mex67-AA 25 °C chase datasets using a linear and exponential model compared to published datasets (Miller et al., 2011; Sun et al., 2012; Neymotin et al., 2014; Presnyak et al. 2015; Chan et al., 2018). **H**. Scatterplot shows the correlation of mRNA half-lives calculated from the Mex67-AA 25 °C chase datasets using a multiple fit linear model (x-axis) or an exponential model (y-axis). Blue lines show the median half-lives in each dataset.

Single gene examples (Figure 1C and Supplemental Figures 1B-G) revealed that various transcripts display different decay patterns and that pA-tail shortening co-occurs with transcript degradation. While mRNA decay is easily represented as a reduction in mRNA abundance, polyadenosine tail length distribution in control cells is usually normal, representing mRNAs at various stages of deadenylation (Figure 1D). This makes modeling the latter process difficult. Therefore in our collection of chase datasets, we described changes in pA-tail lengths distribution using following quantile values: 95^th^, 90^th^, 85^th^, 80^th^, 75^th^, 50^th^, 15^th^, 10^th^ and 5^th^ (Figure 1D). Displaying the changes of each quantile value as a fraction of the origin revealed that data could be fitted into a line in a similar manner as mRNA decay (Figure 1E and Supplemental Figures 1B-G).

The absence of cytoplasmic polyA polymerases in budding yeast guarantees that upon mRNA synthesis arrest, the pA-tails can only be shortened or remain stable over time. This implies that the potential changes in pA-tail lengths within the group of mRNAs with the longest pA-tails, which should cluster newly-made mRNAs, should reveal ‘pure’ deadenylation dynamics, while mRNAs with the shortest pA-tails are potentially less indicative of deadenylation rates, as they are more likely to include fractions of mRNAs with stable pA-tail-lengths overtime. To construct our model, we thus used only the values of the upper quantile (95-75^th^) since inspection of several examples (Figure 1D and Supplemental Figures 1B-G) suggested that the pA-tail lengths of the upper quantiles represent lengths close to newly made mRNA polyadenosine tails (Turtola et al, 2021), while the 50^th^ quantile is closer to the mean and median values, more likely representing transcripts deadenylated prior to the commencement of the experiments. Due to low levels of transcripts in the latter time points, which caused inaccurate pA-tail estimations only time points up to 20 min were used to model deadenylation (Supplemental Figure 1A).

Examination of the bulk mRNA abundance and pA-tail length change by quantiles (Supplemental Figure 2A-J) prompted us to test both a linear and an exponential model to describe the data accurately. The linear model was built by linear regression or a multiple linear fit as described in materials and methods. We found that bulk mRNA half-lives derived from the exponential and linear models equaled 15.2 and 13.5 min, respectively (Figure 1G), and were within the range expected from the literature, displaying a high to moderate Spearman correlation (0.78 to 0.41) with published datasets obtained using various techniques (Figure 1H; Miller et al., 2011; Sun et al., 2012; Neymotin et al., 2014; Presnyak et al. 2015; Chan et al., 2018). Function parameters produced using each model were well-fitted to experimental data for single mRNA examples (Figure 1D, 1F and Supplemental Figures 1A-F). The linear and exponential half-life estimates from the export-block chase strongly agreed for both models (Spearman rho = 0.98), diverging only within the range of marginally short half-lived mRNAs (Figure 1I) and non-coding RNAs (Supplemental Figure L) for which the number of time points collected was not sufficient. This validates our experimental approach to modeling of mRNA decay. Henceforth, the decay and deadenylation rates will be characterized by the slope factor value or the decay factor describing respectively the linear or exponential functions fitted into the data (listed in Supplemental Table 1).

### Deadenylation and decay rates correlate positively transcriptome-wide and are linked to transcript function and codon optimality

To define the putative causative relationship between deadenylation and decay, we compared function parameters calculated from our models and found a strong correlation between both equal to 0.72 in the exponential and 0.59 to 0.68 in the linear regression and multiple linear fit models, respectively (Spearman rho; Figure 2A-B and Supplemental Figure 2P). This suggests a causative relationship between deadenylation and decay, as was previously suggested (De Magistris, 2021).

**Figure 2.**
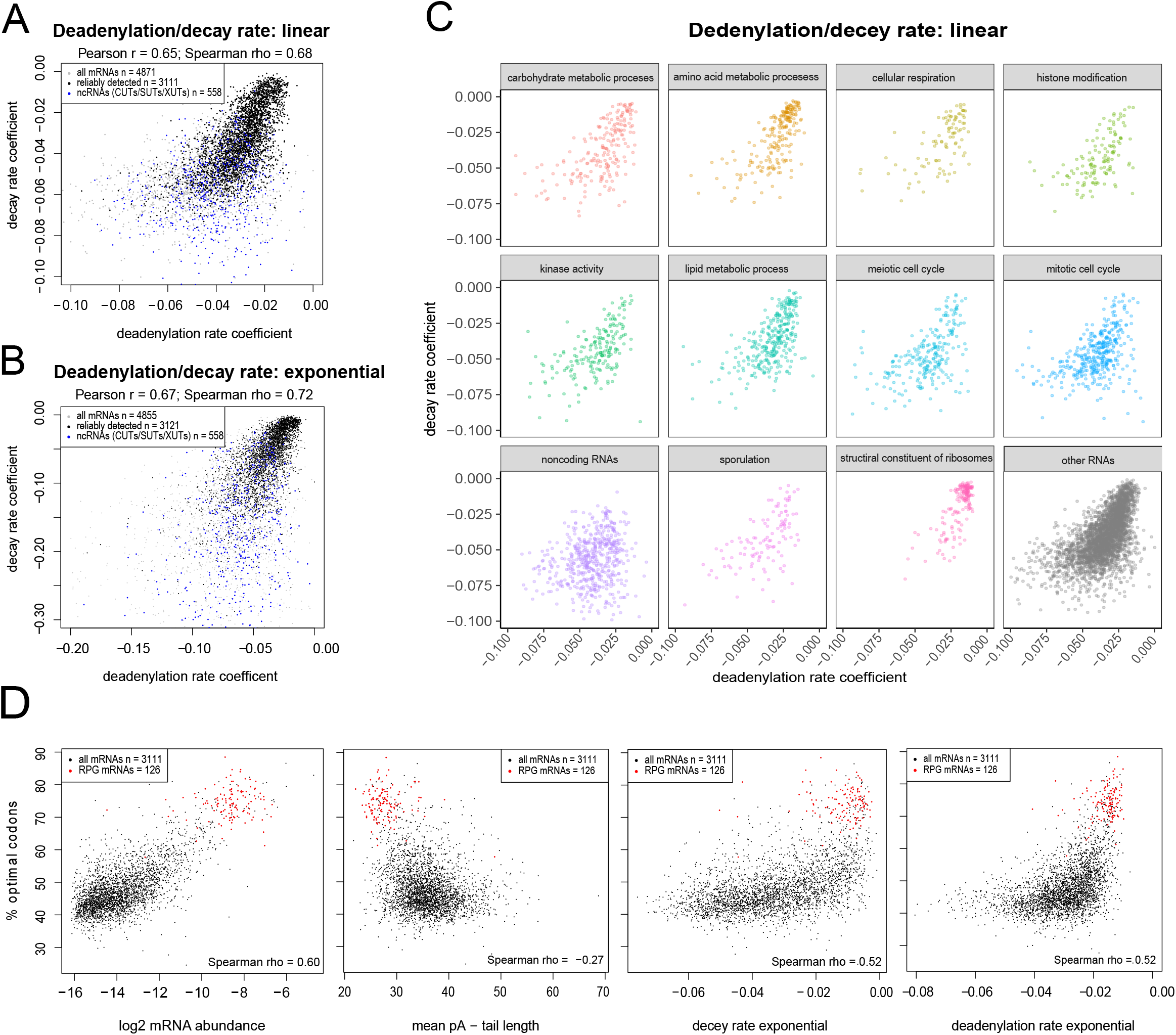
Decay and deadenylation rates correlate at steady-state and are linked to codon optimality. **A-B**. Scatterplot shows the transcriptome-wide relationship between the deadenylation and decay rates derived from multiple fit linear (A) or exponential (B) models. **C**. Scatterplot matrix of the plot showed in displaying deadenylation and decay rates of various mRNA groups clustered by gene-ontology function. **D**. Scatterplot matrix showing on the y-axis the percent of optimal codons and on the x-axis mRNA abundance (top-left panel), mean pA-tail length (top-right panel), linear decay rate (bottom-left panel) and linear deadenylation rate (bottom-right panel).

Deadenylation and decay rates of mRNAs were on average slower than non-coding RNAs (Figure 2A-B), consistent with the rapid removal of those mostly non-functional transcripts. This prompted us to further examine deadenylation and decay rates of several coding transcript gene-ontology clusters (Figure 2C). We found that evolutionarily conserved groups of transcripts encoding for ribosomal protein subunits (Ribosomal Protein Gene mRNAs, henceforth called RPG mRNAs) and factors responsible for amino-acid metabolic process cluster together and are characterized by slow deadenylation and decay. In contrast, genes regulating mitosis formed a much less compacted cluster of mRNAs with short half-lives and a fast deadenylation rate. This is consistent with the notion that the production of mitotic regulators is only required by the cell during a short period of time. Deadenylation and decay rates of other functional groups of transcripts were scattered across the entire distribution. Note that there was a strong correlation between mRNA abundance and either decay or deadenylation rates (Supplemental Figure 2K). This again confirms that mRNAs of high expression, which are mainly house-keeping, are deadenylated and degraded slowly in a pattern close to a linear model. Since mRNA abundance and mean pA-tail length are anti-correlated (Supplemental Figure O; Tudek et al., 2021), the weaker connection between decay and deadenylation rates and pA-tail length is likely a secondary correlation.

Previous works suggested translation rates define mRNA degradation speed, with transcripts bearing optimal codons being degraded slowly and those with rare codons displaying the shortest half-lives (Presnyak et al., 2015; Harigaya et al., 2016; Radhakrishnan et al., 2016). Codon optimality and mRNA abundance showed a 0.6 positive Spearman correlation (Figure 2D first panel). This means that on a transcriptomic level, conserved, highly abundant, and mostly housekeeping genes have the highest percentage of optimal codons and cluster separately from most of the transcriptome. RPGs mRNAs stand out with the most optimal codons. Though a similar bi-modal distribution can be seen for codon optimality and mean pA-tail length, both factors were not correlated (Figure 2D, second panel). Remarkably, both deadenylation and decay rates revealed a 0.5 Spearman correlation with codon optimality and together displayed an exponential distribution (Figure 2D third and fourth panel and Supplemental Figure 2R). Effectively, this shows that codon optimality is a feature that delays the deadenylation and decay of a relatively small percentage of abundant mRNAs. To further confirm that, we cross-compared our data to the model produced by Siwiak and Zielenkiewicz (2010) and observed that decay and deadenylation rates were much less correlated with ribosome density than they were with the time required to translate a single mRNA (Supplemental Figure 2R), confirming the predicted notion that slower translation through rare codons can accelerate decay (Presnyak et al., 2015; Radhakrishnan et al., 2016; Webster et al., 2018) and also deadenylation.

### Deadenlyation is the fastest on newly made pA-tails and linked to Pab1 binding

Our deadenylation models are built on the pA-tail length changes in the upper quantiles of the pA-tail distribution. The 75^th^ – 95^th^ quantiles largely comprised pA-tails containing between 30 and 80 adenosines, which – based on previous Pab1 footprint experiments (Figure 3A; Webster et al., 2018; Baer and Kornberg, 1980; Schäfer et al., 2019) – should bind from two to three copies of this major cytoplasmic polyA-binding protein and correspond to pA-tail lengths of newly produced mRNAs (Turtola et al., 2021; Tudek et al., 2021). In contrast, the mean/median mRNA pA-tail lengths and the 50^th^ quantile are consistent with values corresponding to one or two Pab1 molecules (Figure 3A). We observed that the deadenylation rates calculated separately across each quantile tend to decrease for lower quantiles (Figure 3B, and Supplemental Figure 3A), and hypothesized that it was due to Pab1-mediated regulation of mRNA deadenylation. To test the *in vivo* role of Pab1 in deadenylation, we depleted this protein using an auxin-inducible degron (AID) system for 1 and 2 hours (Supplemental Figure 3B; Nishimura et al., 2009; Morawska and Ulrich, 2013). Pab1 depletion resulted in time-dependent elongation of mean mRNA pA-tail lengths (Figure 3C and Supplemental Figure 3C). Examination of median quantile distributions revealed no change in the upper 95^th^ or lower 5^th^ quantile values, but a marked increase in all intermediate quantiles (Supplemental Figure 3D). Thus, loss of Pab1 does not significantly alter the newly made pA-tail lengths but rather delays deadenylation. Altogether, in agreement with previous *in vitro* (Webster et al., 2018) and *in vivo* (Sachs & Davis, 1989) observations, our results show that Pab1 strongly stimulates deadenylation on newly made long pA-tails, which bear 2 to 3 Pab1 copies (Schäfer et al. 2019), but can become a factor limiting deadenylation speed when the pA-tail is shorter and bound by only one Pab1 molecule.

**Figure 3.**
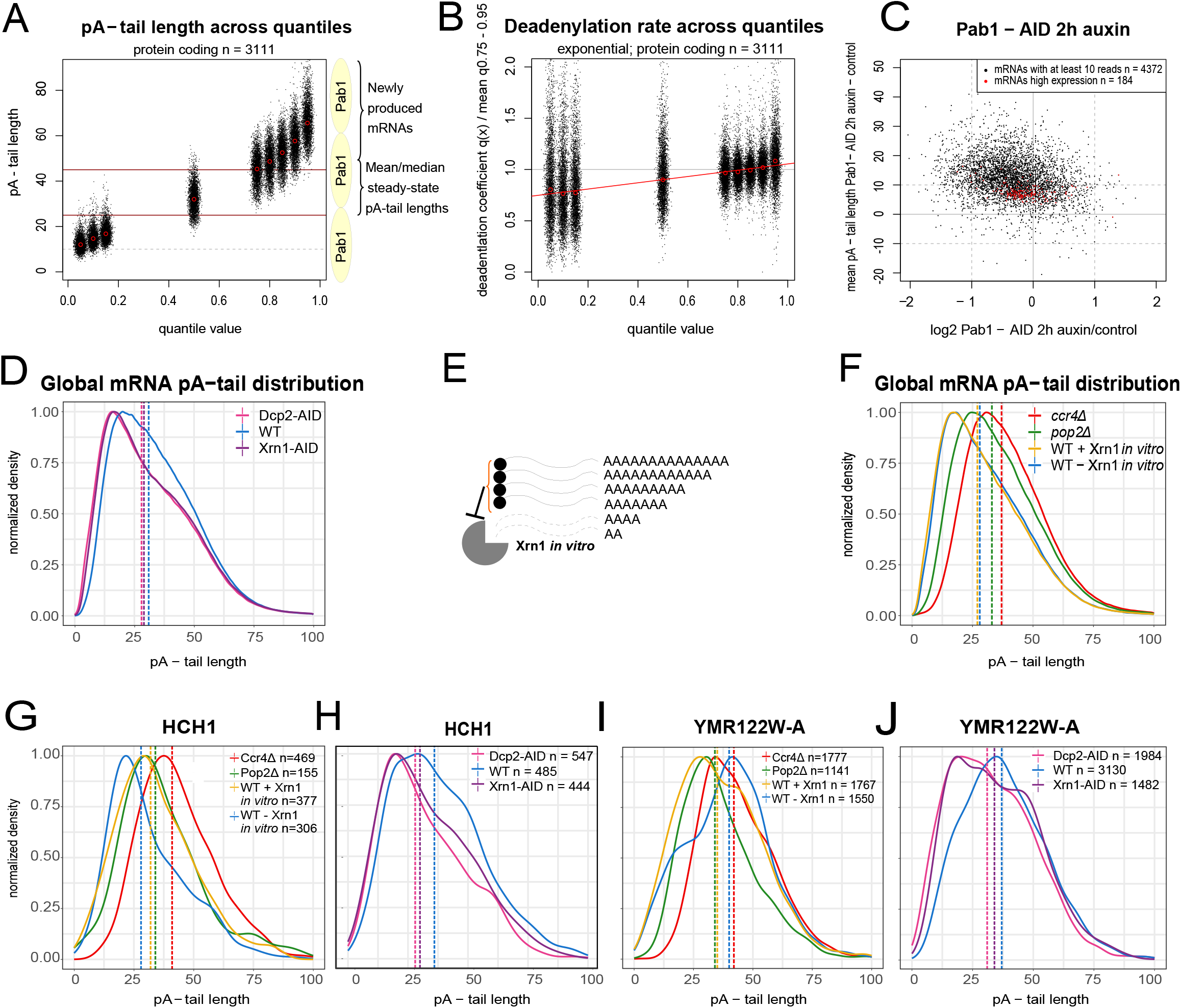
Pab1 is required for *in vivo* deadenylation and decapping commonly occurs at steady-state mRNA pA-tail lengths. **A**. Scatterplot shows the distribution of pA-tail quantile values in wild-type cells. On the right side of the graph, schematic representations of the Pab1 polyA-binding protein are shown to outline that the region bound by one molecule is around 22-30 As. **B**. Scatterplot shows the quantile-specific deadenylation rate estimate calculated using the exponential model relative to the transcript deadenylation rate mean obtained by averaging deadenylation rates of the upper quantiles (75^th^ – 95^th^). **C**. Scatterplot shows the change in mRNA abundance on the x-axis relative to the absolute change in the mean pA-tail length of mRNAs upon 2 hours of Pab1 depletion using the AID system. **D**. Graph shows the global pA-tail length distribution of coding transcripts in control cells compared to Dcp2 and Xrn1 depletion using AID. The distribution shows all pA-tail lengths regardless of transcript abundance, thus mRNAs of high expression are overrepresented. **E**. Scheme shows that Xrn1 can only digest mRNAs devoid of a cap, potentially altering pA-tail distribution if transcripts are only decapped at a defined pA-tail length. **F**. Graph shows the global pA-tail length distribution of coding transcripts in control cells in comparison to wild-type sample digested *in vitro* with Xrn1 and in *pop2Δ*. **G-H**. Graphs show pA-tail length distributions of *HCH1* mRNA in (G) control cells compared to wild-type samples digested *in vitro* with Xrn1 and in *pop2Δ* and for (H) control cells compared to Dcp2 and Xrn1 depletion using AID. **I-J**. Same as (G-H) for *YMR122W-A* mRNA.

### Decapping is initiated at 20-35 adenosine-long pA-tails

Decapping was suggested to be initiated when the pA-tail is short (Chen and Shyu, 2011; Decker and Parker 1993), but the exact pA-tail length at which degradation is initiated has not been determined yet. To clarify this relationship between pA-tail length and decapping, we first depleted the decapping enzyme Dcp2 and the Xrn1 5’-3’ exonuclease for 2 hours using the AID system (Supplemental Figures 3E, 3F). The global mRNA pA-tail profile revealed a clear shift towards shorter pA-tails (Figure 3D), which was validated on a bulk mRNA distribution profile (Supplemental Figure 3G). Accordingly, at the single mRNA level, loss of both factors resulted in a change of the most abundant median pA-tail from 31-35 As in control cells to 26-30 As in Dcp2-or Xrn1-depleted cells (Supplemental Figure 3H). Those phenotypes could be due to the stabilization of short-tailed mRNAs, otherwise degraded in a wild-type situation. Alternatively, infrequently produced transcripts with short tails could accumulate because the half-life of mRNA is extended by inhibition of decay, allowing more time for deadenylation. The latter hypothesis is supported by the fact that depletion of Xrn1 and Dcp2 results in opposite effects on transcriptomic mRNA levels, with changes in pA-tail length best correlating only within mRNAs of high expression (Supplemental Figure 3I-K), indicating that depletion of these factors leads to indirect effects.

To discriminate between these possibilities, we decided to treat the wild-type RNA sample with Xrn1 *in vitro*, which should digest mRNAs devoid of caps (Figure 3E), in order to assess a more physiological situation. This did not substantially alter the global pA-tail length profile or mRNA abundance (Figure 3F, Supplemental Figure 3L), consistent with the notion that decapped mRNAs are not stable and only scarcely present in a wild-type strain. However, inspection of single mRNA pA-tail profiles showed that in many cases, the fraction sensitive to Xrn1 *in vitro* digestion was slightly shorter than the mean, and also stabilized in Dcp2 and Xrn1 AID-depleted cells (Figure 3G, 3H and Supplemental Figures 3M, 3N). At times, a long pA-tailed fraction was clearly subject to decapping (Figure 3I, 3J and Supplemental Figures 3O, 3P). Overall, these results suggest that decapping is initiated close to the individual mRNA median pA-tail length, that is, between pA-tail lengths of 20-35 As. This is clearly longer than the estimation for mammalian cells (Eisen et al., 2020).

Surprisingly, a wild-type RNA profile digested with Xrn1 was at times similar to the one found in *pop2Δ* cells (Figure 3G-J and Supplemental Figures 3M-P), with changes in mean pA-tail length tending to correlate (Supplemental Figure 3R). A Pop2 *in vivo* impact on deadenylation was suggested to be less prominent than that of Ccr4 (Tucker et al., 2002; Webster et al., 2018). Though loss of *POP2* resulted in a weak change in bulk mRNA pA-tail length distribution compared to *ccr4Δ*, (Figure 3E), changes in both *pop2Δ* and *ccr4Δ* relative to the wild-type are highly correlated (Supplemental Figure 3S). Altogether, those results might indicate that the final steps of deadenylation are often mediated concurrently to decapping and occur near steady-state median pA-tail lengths, when the tail is still associated with one Pab1 molecule.

### During heat-stress response decay rate increases more dramatically than the deadenylation speed

To further investigate the potentially causative relationship between deadenylation and decay, we turned to stress conditions. Previous works showed that during acute response to elevated temperature, a couple of hundred coding loci are transiently transcriptionally silenced (Vinayachandran et al., 2018). We produced three replicates of a heat-stress chase experiment. In parallel, we also sequenced two replicates of a chase experiment using thiolutin at 25 °C (Supplemental Figure 4A). The drug is cited as a general transcriptional inhibitor, but some works show that it selectively induces transcription of genes related to the stress response (Adams and Gross, 1991; Neymotin et al. 2014), and might thus mimic some of the processes observed during adaptation to growth at 37 °C. Indeed, in cells treated with thiolutin, we observed up-regulation of mRNAs transcriptionally induced in response to heat-shock, but to a lower degree than in cells directly subjected to heat-shock (Figure 4A and Supplemental Figure 4B). This suggests that a milder stress response is induced following drug treatment.

**Figure 4.**
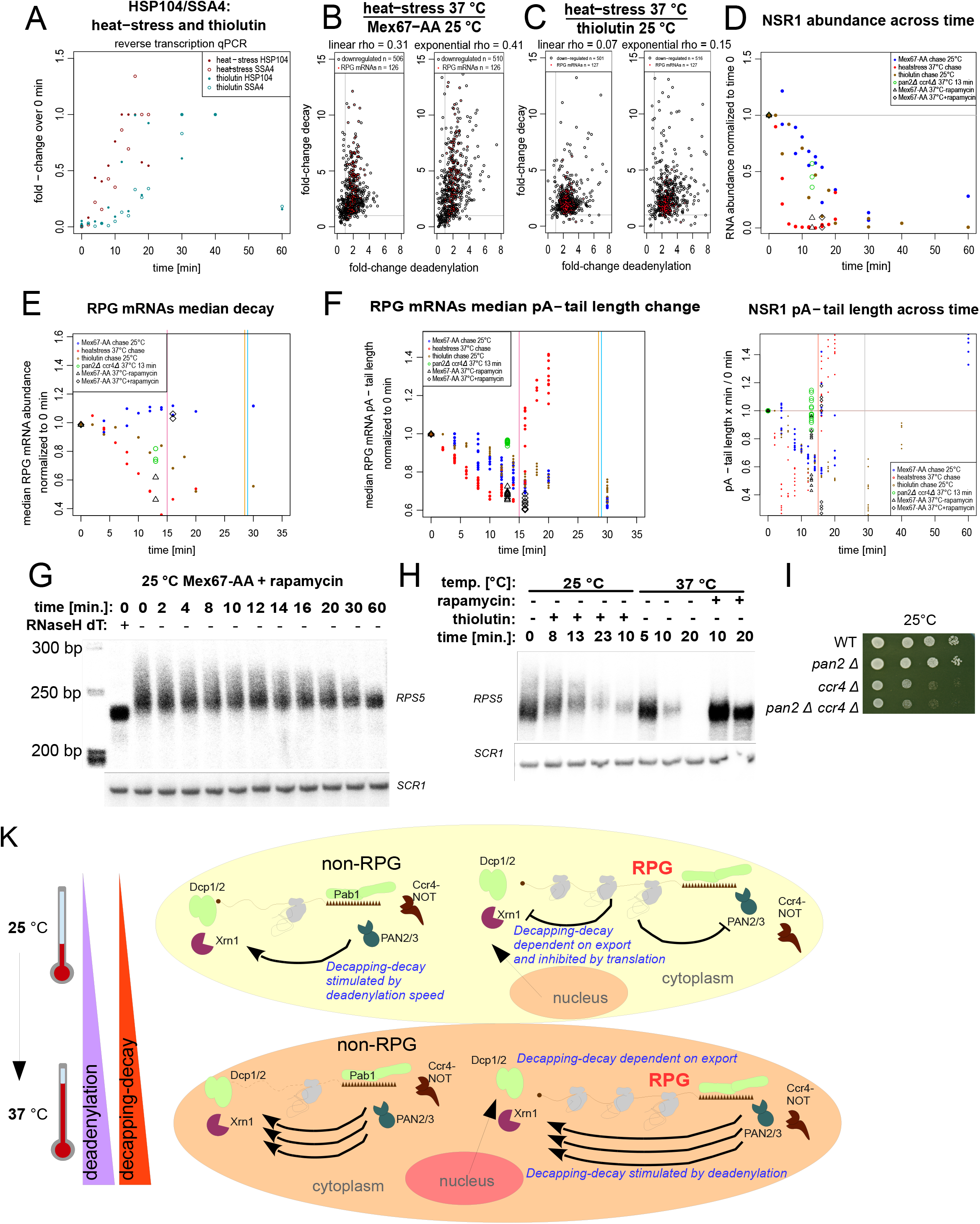
Deadenylation accelerates but is not a prerequisite for mRNA decay. **A**. Graph shows the up-regulation of *HSP104* and *SSA4* mRNA levels during heat stress and thiolutin treatment at 25 °C by reverse transcription coupled to qPCR. **B-C**. Comparison of deadenylation and decay rate changes for mRNAs transcriptionally silent during heat stress (Vinayachandran et al., 2018) for (B) heat-stress compared to Mex67-AA 25 °C and (C) heat stress compared to thiolutin treatment 25 °C. **D-E**. Decay (D) and (E) deadenylation profile for a non-RPG *NRS1*. **F**. Graph shows the median decay curve of RPG mRNAs in Mex67-AA 25 °C, thiolutin 25 °C and heat stress chase datasets compared to heat stress single time points for the double *ccr4Δ pan2Δ* mutant and Mex67-AA cells export-blocked or control. To block nuclear export, rapamycin was added 3 min prior to heat-stress and thus those time points are plotted 3 min later (16 min) relative to the control cells (13 min). The bulk decay curves are shown in Supplemental Figure 4. **G**. Graph shows median deadenylation curve of RPG mRNAs complementary to the decay curve in Figure 3B. The vertical lines delineate the time points, which were not used to calculate deadenylation rates. The heat-stress chase cut-off line was inserted earlier as the stress response is no longer than 15 min and any later point would include *de novo* synthesized mRNAs, as indicated by the increase in upper quantile values. The bulk deadenylation curves are shown in Supplemental Figure 4. **H-I**. Northern blots showing *RPS5* decay and deadenylation in Mex67-AA 25 °C, thiolutin 25 °C, and heat stress chase. *SCR1* non-coding RNA is shown as a loading control. **J**. Growth test comparing wild-type cells to *ccr4Δ, pan2Δ*, and the double *ccr4Δ pan2Δ* mutant at 25 °C. Other temperatures are shown in Supplemental Figure 4. **K**. Model describing the mechanism of mRNA deadenylation and decay. For details see discussion.

We used the heat-stress and thiolutin chase datasets to model deadenylation and decay in the same way as previously for Mex67-AA export-block datasets (Figure 1; Supplemental Figure 4A). We next compared the fold-change in function parameters between datasets (Figure 4B, 4C). Deadenylation and decay rates increased for most of the transcripts under heat stress compared to the Mex67-AA 25 °C chase. However, deadenylation rates increased much less dramatically than decay rates, especially for RPG mRNAs, producing a weak correlation (Figure 4B). Importantly, comparison of both factors in heat stressed cells to those treated with thiolutin at 25 °C yielded no correlation, although both coefficients were roughly two-fold increased in response to elevated temperature (Figure 4C). On a single mRNA level, we found mRNAs for which deadenylation and decay were both accelerated during heat-stress to various extents (*NSR1* - Figure 4D and *NOP1, YEF3* and *EGD1* - Supplemental Figure 4C, D, E). Surprisingly, we also found a collection of transcripts that displayed equal or slightly increased decay rates but were virtually not deadenylated during heat-stress (Supplemental Figure 4F, G, H, I). Those results might support the hypothesis that deadenylation rate is a factor required to increase the speed of decay rather than being indispensable to trigger decapping.

### Deadenylation is not required for the degradation of ribosomal protein mRNAs but accelerates this process under heat-stress

RPG mRNAs were the largest functional group of transcripts down-regulated during heat-stress. Since we previously observed that they displayed similar decay and deadenylation rates (Figure 2C) we next examined them as a group (Figure 2E, 2F and Supplemental Figure 5A-C). During the export block at 25 °C, degradation of most RPG mRNAs was not be observed. In contrast, median half-lives of RPG mRNAs upon thiolutin treatment were 20 min long. The decay of those mRNAs was even faster under heat shock conditions with a median half-life of 12 min. A similar change in RPG mRNA half-lives was observed in variety of previously published datasets obtained using orthogonal methods (Supplemental Figure 5D; Miller et al., 2011; Sun et al., 2012; Neymotin et al., 2014; Presnyak et al., 2015).

Despite marked differences in stability, the deadenylation rates under both thiolutin and export-block conditions were equal. Only elevated heat resulted in accelerated mRNA deadenylation (Figure 2E, 2F and Supplemental Figure 5A-C). Those data were puzzling since if deadenylation rates were causative to decay in all cases, we would expect both factors to change proportionally. Such a relation, however, is not observed when comparing the export-block to the thiolutin treatment. We thus hypothesized that at 25 °C decapping of RPG mRNAs is slow and dependent on a factor that is exported from the nucleus to the cytoplasm, which possibly implies that thiolutin treatment at 25 °C reveals the true half-lives of RPG mRNAs. To test this hypothesis, we blocked export by depleting Mex67 for 3 min before heat stress and observed that decay of the majority of RPG mRNAs was completely inhibited (Figure 2E, 2F; Supplemental Figure 5E, 5F) although selected RPG mRNA did not require export for decay (Supplemental Figure 5G; e.g. RPL4A). At the same time, deadenylation of RPG mRNAs was accelerated for Mex67-AA control and export-blocked cells as it was in wild-type cells subjected to heat stress (Figures 4E-F, black triangles and squares; Supplemental Figure 4F; single gene examples – Supplemental Figure 6). This was validated by reverse-transcription coupled to qPCR and northern blotting analysis (Figure 4G, 4H; Supplemental Figures 5F). This shows that deadenylation rate is completely independent of decay speed.

Altogether, our data indicate that at steady-state, RPG mRNA decay is dependent primarily on an unknown factor exported from the nucleus. However, faster deadenylation is expected to accelerate the degradation of those transcripts during heat-stress. To test this, we heat-stressed a double *ccr4Δ pan2Δ* mutant, in which, as expected, deadenylation was almost completely blocked (Figure 4E and F and Supplemental Figure 3H). Still, RPG mRNAs were degraded with a decay curve strikingly similar to one of wild-type cells treated with thiolutin at 25 °C (Figure 4E and Supplemental Figure 5H). A decrease in the decay of RPG mRNAs was also observed in single *ccr4Δ* and *pan2Δ* mutants (Supplementary Figure 5I), suggesting that both deadenylases contribute independently to accelerated decay. In agreement with Bresson et al., (2020) decay of RPG mRNAs was dependent on decapping by Dcp2 (Supplemental Figures 5F) and supported by decapping complex co-factors *dhh1Δ* and *lsm1Δ* (Supplementary Figure 5I). Thus deadenylation boosts RPG mRNA decay under stress but is not required for decapping and degradation to occur. The fact that decay is not completely inhibited when deadenylation is blocked, explains the viability of the double *ccr4Δ pan2Δ* mutant (Figure 4I and Supplemental Figure 5J). Importantly, the exact same relations between deadenylation and decay were observed on a individual transcript level (Supplemental Figure 6A-D). Altogether we show for the first time that decapping can be uncoupled from deadenylation for a large group of transcripts.

## DISCUSSION

In our previous work, we showed that newly made mRNA pA-tails are typically 60 adenosines long and that this length is defined by Nab2 nuclear polyA-binding protein (Turtola et al., 2021; Tudek et al., 2021). At steady-state, most mRNAs display a normal pA-tail length distribution, in which the most abundant adenosine tail often matches the region required for binding of one copy of cytoplasmic polyA-binding protein Pab1 (22 to 30 As; Webster et al., 2018; Subtelny et al., 2014; Tudek et al., 2021). This indicates that it is the most stable cytoplasmic form of the mRNA. Consistently, deadenylation is more rapid when the pA-tail is long, newly made, and bound by multiple Pab1 subunits. From this, the most direct interpretation is that in yeast, the primary role of deadenylation is to form a stable form of the transcript, bound by one Pab1 molecule, on which deadenylation will occur relatively slowly (Figure 4K). Such interpretation is consistent with the closed-loop model that exploits the interaction between Pab1 poly(A)-binding protein and the translation initiation cap binding complex (Tarun et al., 1997; Otero et al., 1999; Archer et al., 2015). The redundant action of both PAN2/3 and Ccr4-NOT deadenylases mediates initial deadenylation in a manner dependent on the polyA-binding protein. In accordance with this hypothesis, previous works have shown that Pab1 *in vitro* binds to both PAN2/3 and Ccr4-NOT (Webster et al., 2018; Schäfer et al., 2019; Wolf et al., 2014).

Using two alternative approaches, we found that decapping is often initiated close to the most prevalent pA-tail length, between 20 and 35A, and weakly correlates with additional deadenylation (Figure 5). This estimate is seemingly longer than the 25 adenosines previously settled by modeling of deadenylation and decay in mouse cell lines (Eisen et al., 2020), though both studies agree on the notion that decapping occurs when the pA-tail can still be bound by one polyA-binding protein.

On a transcriptomic level at steady-state conditions, decay and deadenylation are strongly positively correlated, seemingly suggesting causation, in accordance with early models which proposed deadenylation was a prerequisite for decay (Chen and Shyu, 2011; Yi et al., 2018; Decker and Parker, 1993). Though deadenylation and decay rates at steady state display a continuous distribution and are correlated with transcript abundance, gene-ontology analyses show that some groups of transcripts tend to have a similar deadenylation and decay speed. This indicates a functional organization of the transcriptome metabolism exists. Transcripts responsible for the regulation of cell division are expressed in low levels and have a relatively short half-life, reflecting their transient role in the cell. In contrast, mRNAs encoding proteins controlling basic cellular processes, such as amino acid metabolism and RPG mRNAs, are abundant, stable, and slowly deadenylated. In the case of RPG mRNAs, this coordinated regulation also occurs during cellular response to stress, when both deadenylation and decay are accelerated. RPG transcripts are conserved and old, ant this is reflected by the particularly high codon optimality score. The connection between codon optimality and transcript decay speed was shown mostly in reporter transcripts (Presnyak et al., 2015). We, however, show that decay and deadenylation rates are tied to codon optimality in an exponential manner. Effectively this sets aside mRNA of basic cellular metabolism, such as RPGs, from the rest of the transcriptome but also indicates that minor codon adaptations, which facilitate translation, can potentially have a big impact on transcript stability and protein expression.

Regardless of the general correlation between pA-tail shortening and degradation, several of our observations place deadenylation as a factor auxiliary to decay rather than causative (Figure 4K). First, though accelerated decay under heat-stress conditions coexists with increased deadenylation rates, both factors are poorly correlated, with decay being more strongly affected than deadenylation. Second, in a double *ccr4Δ pan2Δ* mutant, decay under stress conditions is not as fast as in wild-type cells, but not entirely blocked, consistent with the viability of the double mutant. Third, we observed that decay and deadenylation can be uncoupled, as exemplified by some mRNAs during heat-stress, which are decapped, but not deadenylated. Similarly, most RPG mRNAs were slowly deadenylated under steady-state growth conditions, but their ultimate decay was fully dependent on ongoing export. The RPG decay mechanism might rely on a yet unidentified decapping complex co-factor. This co-factor might shuttle from the nucleus and only shortly resides in the cytoplasm or be *de novo* transcribed in the same time, having a short cytoplasmic half-life. We were however unsuccessful in identifying it.

The interesting question is the mechanism of acceleration of decay and deadenylation on most transcripts during heat-stress. This might be due to heat-dependent changes in the kinetics of the complexes involved in both processes. However, the lack of deadenylation of some mRNAs coupled with an unchanged decay rate during heat-stress argues against this. Alternatively, the association of both deadenylases and the decapping complex with their targets might be substantially altered by the activity of supporting co-factors under stress conditions. Considering the number of proteins interacting with deadenylases and Dcp1/2, addressing these possibilities further will require a systematic global approach.

## ACKNOWLEDGEMENTS

This work was mainly supported by National Science Centre Poland, SONATA grant [2020/39/D/NZ2/02174 to A.T.] and by the TEAM/2016-1/3 Foundation for Polish Science grant (to A.D.). A.D. is also a recipient of the ERA Chair’s position, funded by the EU (agreement no. 810425). Work by RT was supported by National Science Centre Poland, SONATA BIS grant [2017/26/E/NZ1/00724] and by National Centre for Research and Development [LIDER/35/46/L-3/11/NCBR/2012]. S.M.’s work was supported by a Polish National Science Centre grant (no. 2020/38/E/NZ2/00372 to S.M.).M.T. is supported by the Academy Research Fellow grant (nos. 349698 and 353682) awarded by the Academy of Finland. Work by M.T. in the laboratory of T.H.J. was supported by a Federation of European Biochemical Societies long-term fellowship and an EMBO long-term fellowship (ALTF 328-2019).

## DATA AVAILABILITY

The accession number for sequencing data are listed in Supplemental Table 4, raw data are deposited at European Nucleotide Archive (ENA) under accession number XXXX. Raw non-sequencing data along with tables containing most relevant statistics for each sequencing dataset are deposited at Mendeley Data at doi: 10.17632/2j3hh37zzs.1.

## CONFLICT OF INTEREST

Authors declare no conflict of interest.

### AUTHOR CONTRIBUTIONS

ACC quality processed the raw data and provided tables with sequencing statistics (mRNA count number, mean, median pA-tail and quantile values), analyzed the median changes in pA-tail length upon Dcp2-/Xrn1-AID depletion or Xrn1 *in vitro* digestion, edited the manuscript. MT produced the Pab1-AID strain, prepared the Pab1-AID samples for sequencing, produced *RPS5*/*SCR1* Northern blots and edited the manuscript. RT tested the bulk pA-tail distribution in Mex67-AA and Dcp2-/Xrn1-AID compared to control, expressed and purified recombinant Xrn1 for *in vitro* digestions, and edited the manuscript. PK supervised raw sequencing data processing. WO and SW synthesized sequencing libraries. US produced the Dcp2-AID strain. THJ acquired funding and edited the manuscript. AD acquired funding and coordinated the research, edited the manuscript. AT conceptualized the project, coordinated the project tasks performed by collaborators, prepared samples for sequencing, analyzed the data, wrote the publication and acquired funding.

## SUPPLEMENTAL FIGURES

**Supplemental Figure 1.**
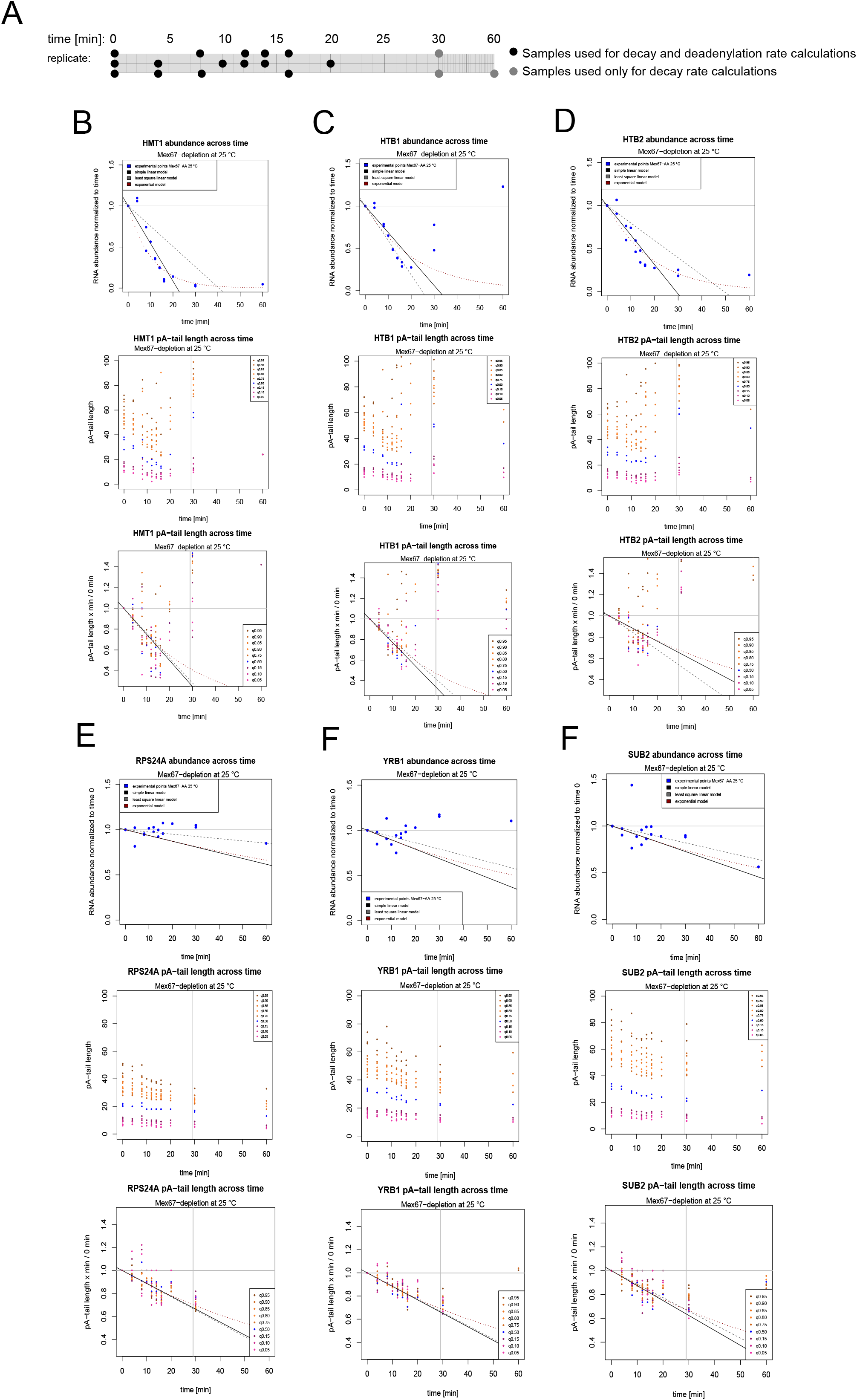
A. Scheme showing the time points collected across three datasets of Mex67-AA chase at 25 °C with an outline of the utility of each point for decay or deadenylation rate calculation. **B**.-**G**. Single transcript examples showing RNA abundance (top panel), raw (middle panel) and normalized (bottom panel) pA-tail quantile distribution change across time in three Mex67-AA 25 °C chase datasets combined.

**Supplemental Figure 2.**
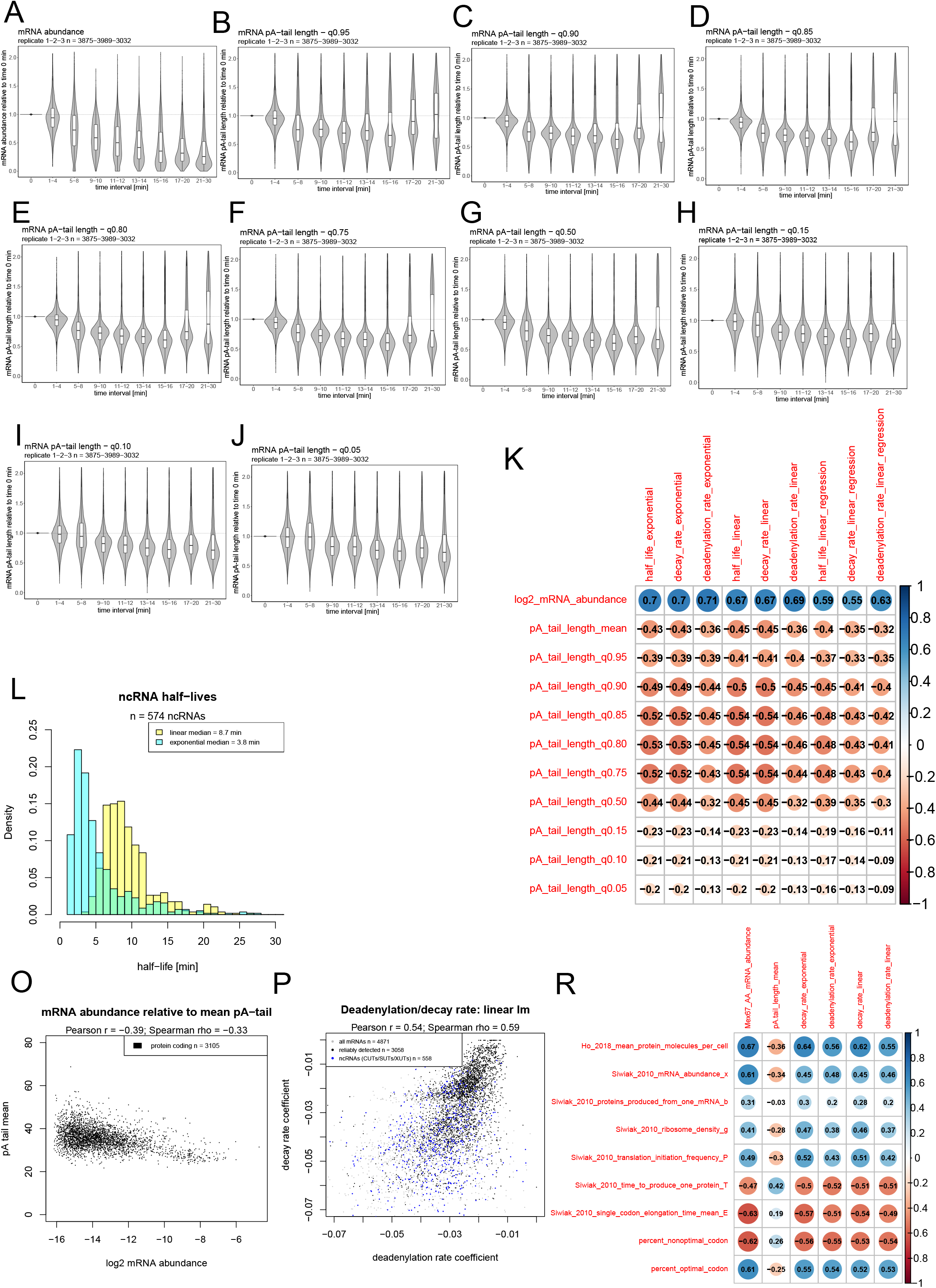
A. Violin plots show change in abundance of over 3 000 coding transcripts reliably detected in three Mex67-AA 25 °C replicates in relation to control time points and grouped in time intervals. **B-J**. Violin plots show change pA-tail quantile (95, 90, 90, 85, 80, 75, 50, 15, 10, 5) values of over 3 000 coding transcripts reliably detected in three Mex67-AA 25 °C replicates in relation to control time points and grouped in time intervals. **K**. Correlation matrix showing the relationship between deadenylation rate, decay rate and half-life calculated by the linear and exponential models and mRNA abundance, mean pA-tail and pA-tail length in each quantile. **L**. Histogram showing distribution of half-lives of ncRNAs (mostly XUTs and SUTs) in the multiple fit linear and exponential models. **O**. Scatterplot showing the relationship between mRNA abundance and mean pA-tail length for Mex67-AA control cells. **P**. Scatterplot shows the relationship between the deadenylation and decay rates derived from a linear regression model. **R**. Correlation matrix between Mex67-AA dataset (containing information about log2 mRNA abundance, mean pA-tail length, and decay/deadenylation rates calculated from the linear and exponential models) and datasets summing Siwiak and Zielenkiewicz (2010) translation model parameters along with Ho et al., 2018 information about mean protein molecules in the cell.

**Supplemental Figure 3.**
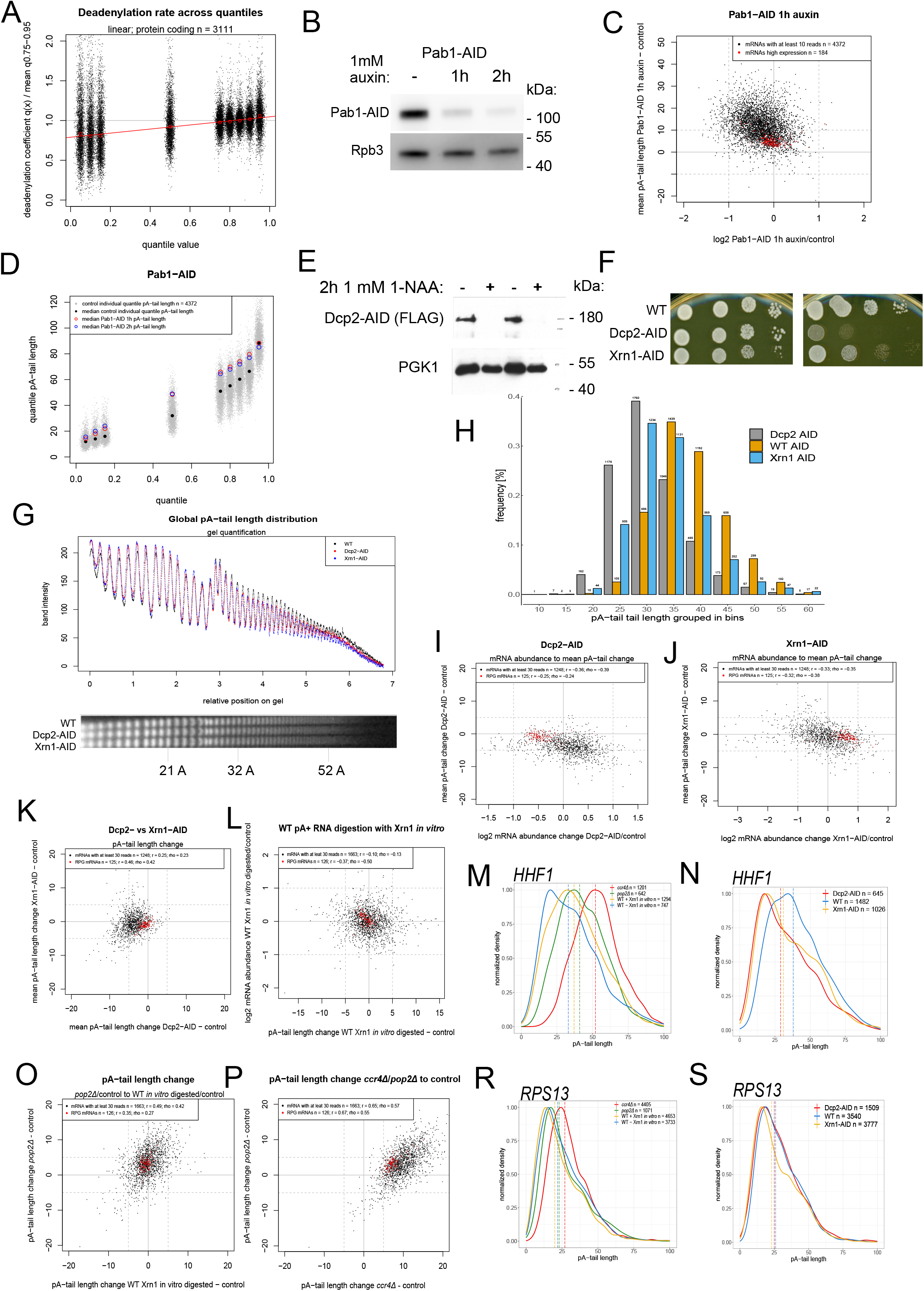
A. Scatterplot shows the quantile-specific deadenylation rate estimate calculated using the multiple fit linear model relative to the transcripts deadenylation rate mean obtained by averaging deadenylation rates of the upper quantiles (75^th^ – 95^th^). **B**. Western blot shows depletion of Pab1 protein using the auxin-inducible degron for 1 and 2 hours. Western blot analysis for Rpb3 protein was used as a loading control. **C**. Scatterplot shows the change in mRNA abundance on the x-axis relative to the absolute change in the mean pA-tail length of mRNAs upon 1 hour Pab1 depletion using the AID system. **D**. Scatterplot shows the distribution of pA-tail quantile values in wild-type cells compared to Pab1-depletion using AID system for 1 and 2 hours. The thin grey dots show individual mRNA pA-tail lengths in control cells, while the larger circles show the median values in both the control (black dot) and Pab1-depleted cells (empty red and blue circles). **E**. Western blot shows the efficiency of depletion of Dcp2 using the AID system for 2 hours. The PGK1 blot is shown as a loading control. **F**. Growth test produced using serial 10-fold dilutions of cells showing depletion of Dcp2 and Xrn1 using the AID system gives the expected growth phenotype. **G**. Bottom panel shows an autoradiogram of the bulk mRNA pA-tail length distributions in control cells and strains depleted for Dcp2 and Xrn1 using the AID system. Top panel shows the quantification of the autoradiogram. **H**. Graph shows the relative number of mRNAs with median pA-tails clustered in bins of 5 for transcripts represented by at least 30 reads for control compared Dcp2- and Xrn1-depleted cells. **I-J**. Scatterplots show the change in mRNA abundance on the x-axis in relation to the absolute change in mean pA-tail length for (I) Dcp2- and (J) Xrn1-depleted cells. **K**. Scatterplot compares the absolute change in mean pA-tail length in Dcp2- and Xrn1-depleted cells. **L**. Scatterplot shows the absolute change in mean pA-tail length in relation to the change in mRNA abundance between a wild-type sample digested or not with Xrn1 *in vitro*. **M-N**. Graphs show pA-tail length distributions of *HHF1* mRNA in (M) control cells in comparison to wild-type sample digested *in vitro* with Xrn1 and in *pop2Δ* and for (N) control cells in comparison to Dcp2 and Xrn1 depletion using AID. **O-P**. Same as (M-N.) for *RPS13* mRNA. **R**. Scatterplot shows the absolute change in mean pA-tail length for wild-type cells treated with Xrn1 *in vitro* compared to control and *pop2Δ* cells in relation to the same control sample. **S**. Scatterplot shows the absolute change in mean pA-tail length for *ccr4Δ* and *pop2Δ* cells compared to the same wild-type sample.

**Supplemental Figure 4.**
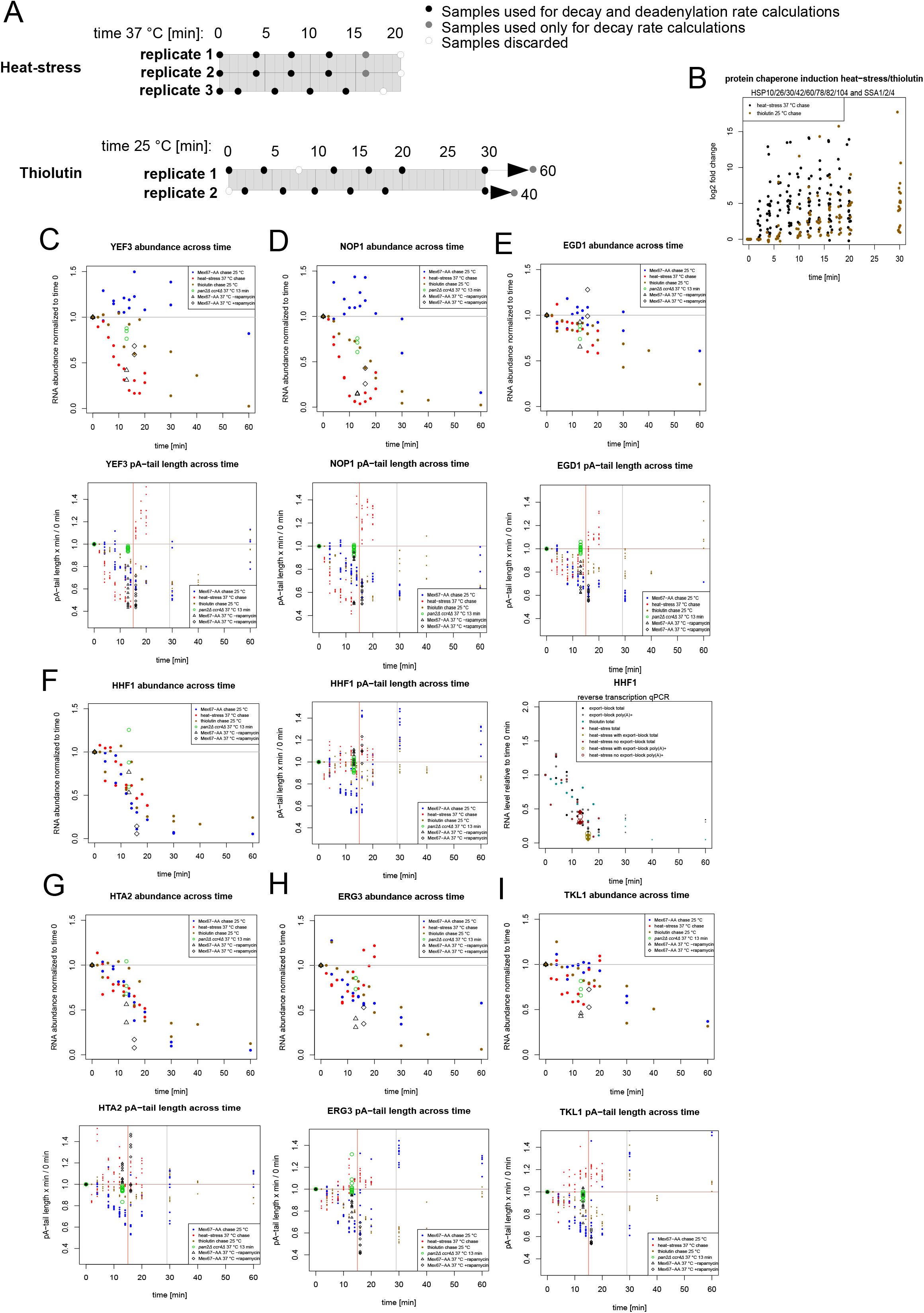
A. Scheme showing the time points collected across three datasets of heat-stress chase at 37 °C and two replicates of thiolutin treatment at 25 °C with an outline of the utility of each point for decay or deadenylation rate calculation. Due to technical reasons, the dataset of thiolutin treatment was normalized to 2 mine time point, which was close enough to the control to salvage the entire set of sequencing. **B**. Scatterplot shows the up-regulation of a collection of chaperone mRNAs in the heat-stress and thiolutin chase sequencing data. Those transcripts are known to be transcriptionally induced during heat stress (Vinayachandran et al., 2018). **C-I**. Graphs displaying decay (top panels) and deadenylation (bottom panels) of single non-RPG mRNAs in the Mex67-AA 25 °C chase, thiolutin treatment at 25 °C, heat stress at 37 °C, heat stress for *ccr4Δ, pan2Δ* mutant and heat stress for export-blocked cells. In case of the *HHF1* mRNA a reverse-transcription coupled to qPCR analysis of the transcript abundance is also shown.

**Supplemental Figure 5.**
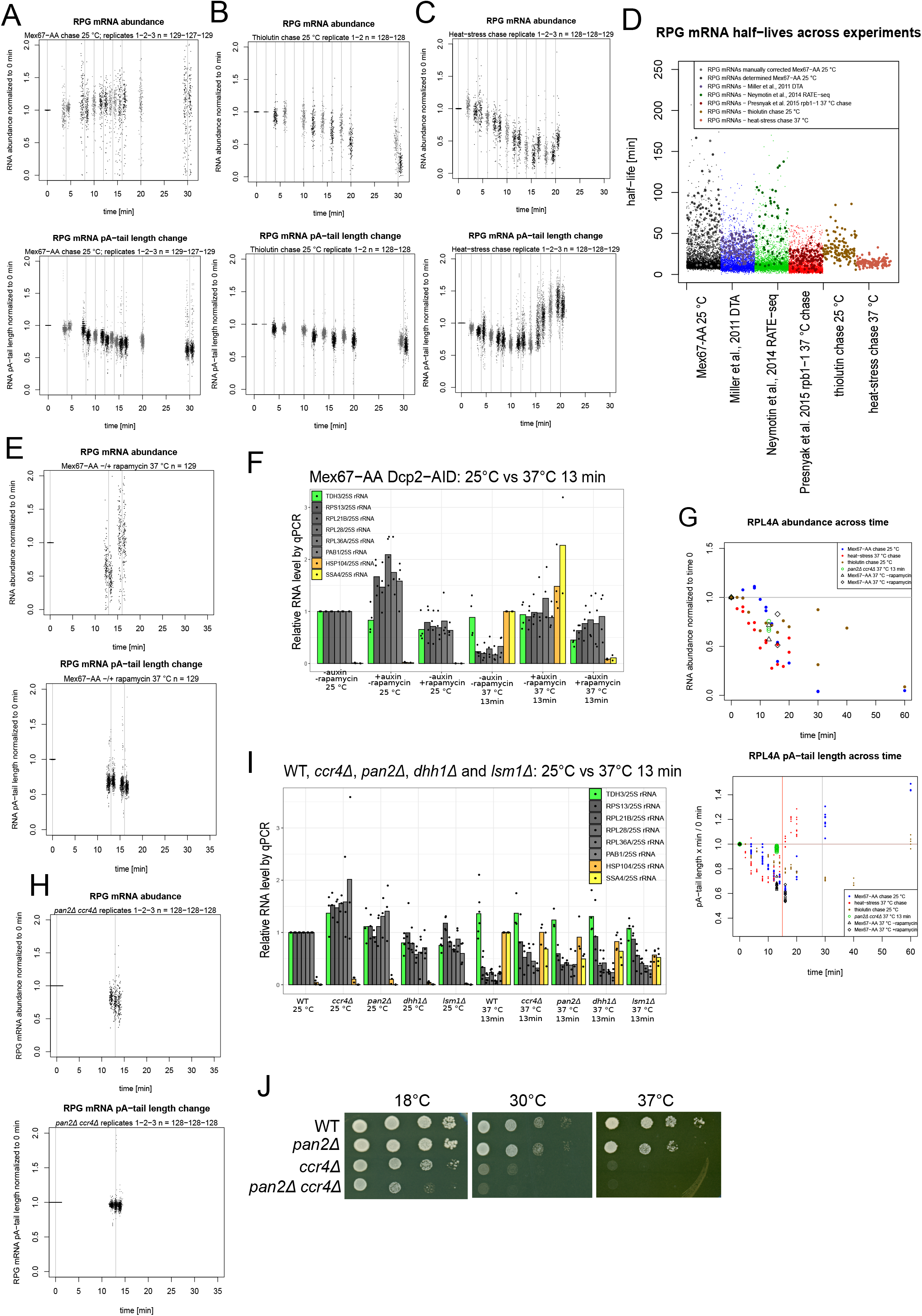
A-C., **E and H** Scatterplots in the top panel show the abundance of RPG mRNAs over time, while those in the bottom panels show the change in pA-tail length in the upper (75^th^ to 95th) quantiles in the following datasets: (A) Mex67-AA 25 °C chase, (B) thiolutin 25 °C treatment, (C) heat-stress 37 °C, (E) Mex67-AA cells export-blocked or not and (H) *ccr4Δ panΔ* heat-stressed cells. **D**. Scatterplot compares the bulk mRNA half-lives in Mex67-AA, thiolutin and heat stress datasets multiple fit linear model to published datasets from (Miller et al., 2011; Sun et al., 2012; Neymotin et al., 2014; Presnyak et al., 2015). The small dots show all mRNAs, while the big ones show the RPG mRNA half-lives. **F**. Barplot shows the abundance of selected RPG mRNAs normalized to 25S rRNA in control or Dcp2- or Mex67-depleted cells at 25 °C compared to 13 min heat shock at 37 °C using reverse transcription coupled to qPCR. Single dots show biological replicate values used to calculate the mean. **G**. Graphs displaying decay (top panel) and deadenylation (bottom panel) of RPL4A in the Mex67-AA 25 °C chase, thiolutin treatment at 25 °C, heat stress at 37 °C, heat stress for *ccr4Δ, pan2Δ* mutant and heat stress for export-blocked cells. **I**. Barplot shows the abundance of selected RPG mRNAs normalized to 25S rRNA in control or *ccr4Δ, pan2Δ, lsm1Δ*, and *dhh1Δ* cells at 25 °C compared to 13 min heat shock at 37 °C using reverse transcription coupled to qPCR. Single dots show biological replicate values used to calculate the mean. **J**. Growth test of comparing wild-type cells to *ccr4Δ, pan2Δ*, and the double *ccr4Δ pan2Δ* mutant at 18, 30, and 37 °C.

**Supplemental Figure 6.**
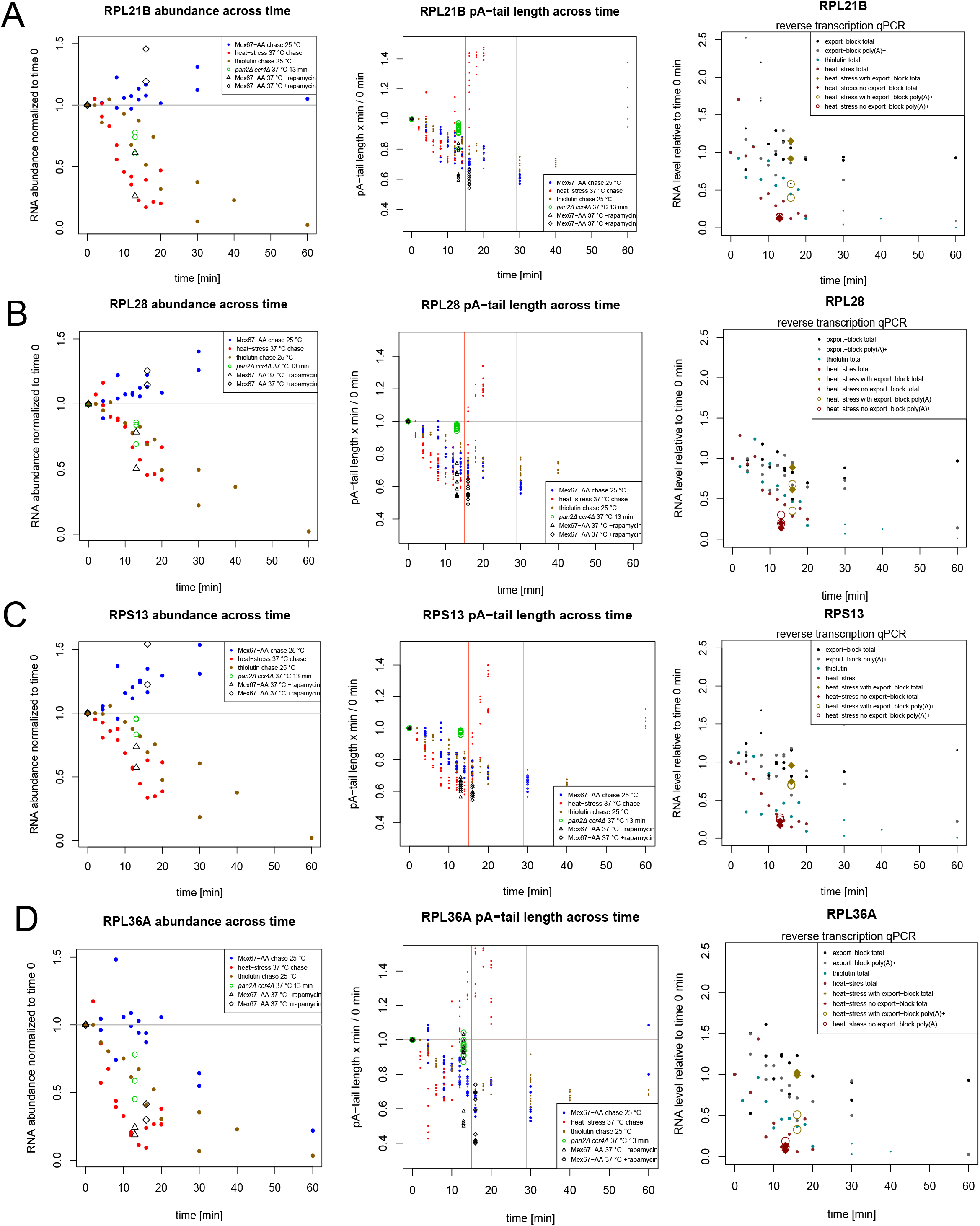
A-D. Graphs displaying decay (left panels) and deadenylation (middle panels) of single RPG mRNAs in the Mex67-AA 25 °C chase, thiolutin treatment at 25 °C, heat stress at 37 °C, heat stress for *ccr4Δ, pan2Δ* mutant and heat stress for export-blocked cells. Right panels for A-D show abundance of those transcripts in the Mex67-AA 25 °C chase, thiolutin treatment at 25 °C and heat stress at 37 °C by reverse transcription coupled to qPCR.

## MATERIALS AND METHODS

### Yeast culture conditions

Yeast cultures were prepared in YPDA media. The Mex67-AA and thiolutin chase experiments were performed at 25 °C. To deplete Mex67 with the Anchor Away tag (Haruki et al., 2008), rapamycin (Cayman Chemicals cat. no. 13346) was added to a final concentration of 1 μg/ml. The heat-stress chase experiment was performed by pre-culturing cells at 25 °C and adding an equal volume of media pre-heated to 51 °C, which resulted in a final temperature of 37 °C. Treatment with thiolutin (Sigma; T3450) was performed by adding the compound to a final concentration of 4 ug/ml. Samples for all chase experiments were collected by mixing the cell culture with an equal volume of ice-cold ethanol, which inactivates cellular metabolism. To deplete Xrn1 and Dcp2 with the AID tags (Nishimura et al., 2009; Morawska and Ulrich, 2013) 1-Naphthaleneacetic acid (1-NAA; N0640-25G, Merck) was added to a final concentration of 1 mM; alternatively, auxin sodium salt (I5148-2G; Merck) was added to a final concentration of 1-3 mM. The list of yeast strains used in this study is shown in Supplemental Table 2.

### Western blot analysis

4-6 OD units of cells were resuspended in 8M urea and incubated at 80 °C. Cells were subsequently mixed with glass beads and disrupted on a vortex for 5 min. The urea cell extract was centrifuged and the supernatant was collected. The protein concentration was measured and after adjusting to equal concentrations extracts were resuspended in Laemmli buffer and 30 μg of protein was loaded on a 6 or 10 % denaturing SDS-PAGE gel. After gel migration proteins were transferred to Amersham nitrocellulose Protran Western blotting membranes (GE10600001) using semi-dry transfer (SEMI DRY BLOT APPARATUS, Bio-Rad, cat. no.1703940). Blots were blocked in TBS-T and 5 % milk and probed with relevant antibodies overnight. Antibodies used were: anti-FLAG (F1804, Sigma), anti-PGK1 (discontinued Novex Life Technology), anti-Pab1 (Santa Cruz Cat. no.: sc57953) or anti-Rpb3 (Abcam 1Y26 cat. no.: ab81859). After washing in TBS-T blots were probed with appropriate secondary antibodies conjugated with HRP in TBS-T 5 % milk for 1 hour, washed, and developed with ECL substrate (Clarity Western ECL, BioRad).

### RNA extraction

RNA was extracted using hot acid phenol method. Cell pellets were resuspended in 400 μl phenol solution saturated with 0.1 M citrate at pH 4.3 (Sigma; P4682) and 400 μl of TES buffer was added (10 mM Tris pH 7.5, 5 mM EDTA, 1 % SDS). Samples were vortexed for 45 min at 65 °C and then centrifuged at 4 °C for 10 min. The supernatant was collected into a fresh tube and 400 μl phenol solution saturated with 0.1 M citrate at pH 4.3 was added. The samples were again vortexed for 20 min at 65 °C and then centrifuged at 4 °C for 10 min. The supernatant was collected into a fresh tube and 400 μl of chloroform was added (C2432; Sigma). The samples were vortexed briefly at room temperature and centrifuged at 4 °C for 10 min. The supernatant was collected into a fresh tube; 45 μl of 2 M LiCl was added and 1 ml of 95 % ethanol. Samples were precipitated for at least 30 min at – 80 °C, centrifuged for 25 min at 4 °C, washed with 400 μl of 80 % ethanol and after removing the supernatant dried at 37 °C. Pellets were resuspended in nuclease-free water and concentration was measured using Nanodrop apparatus.

### Enrichment of the pA^+^ fraction for sequencing library preparation and qPCR analyses

The pA+ fraction was prepared using magnetic beads coupled to oligo-dT from LifeTechnologies (61005). 35 μg of total RNA was resuspended in 50 μl of nuclease free water, if need be a *Schizosacchromyces pombe* total RNA spike-in was added in a w/w ratio of 1:100 to *S. cerevisiae* to provide normalization for sequencing data. RNA was mixed with 50 μl of binding buffer (20 mM Tris-HCl, ph 7.5, 1 M LiCl, 2 mM EDTA) and denatured for 2 min at 65 °C to be cooled on ice. 100 μl of slurry beads per 35 μg of total RNA was pre-washed 3 times in 1 ml of binding buffer and resuspended in 50 μl of binding buffer per sample. Beads were added to the denatured RNA and incubated at room temperature with occasional shaking for 20 min. The supernatant was removed and beads were washed 2 times with 100 μl of wash buffer (10 mM Tris-HCl pH 7.5, 150 mM LiCl, 1 mM EDTA) and after removing any remnants of wash buffer resuspended in 12 μl of nuclease free water. Beads were incubated for 2 min at 80 °C and the supernatant removed from the beads was utilized for library preparation and qPCR analyses as the pA+ fraction.

### Recombinant Xrn1 expression, purification and testing

Nucleotide sequence encoding *Thermothelomyces* (*Myceliophthora*) *thermophilus* Xrn1, with codons optimized for heterologous expression in Escherichia coli, was synthesized by GenScript, provided as a derivative of pUC57 plasmid, and re-cloned into NcoI/XhoI sites of the pET-28b(+) vector (Novagen) using sequence and ligation-independent cloning (SLIC) with primer pair TtXrn1For (5’-ttttgtttaactttaagaaggagatataccATGGGCGTCCCGAAGTTTTTCC-3’) - TtXrn1Rev (5’-atctcagtggtggtggtggtggtgctcgagGCTCTGCAGTGCTGCGGTCTG-3’) for insert amplification in PCR.

Resulting pET-28-TtXrn1-6xHis recombinant vector was introduced into E. coli BL21-CodonPlus(DE3)-RIL chemo-competent cells (Agilent; E. coli B F– ompT hsdS[rB– mB–] dcm+ Tetr gal λ[DE3] endA Hte [argU ileY leuW Camr]) by heat shock-based transformation. Transformants were selected in a standard solid Luria-Broth (LB) medium supplemented with 50 μg/mL kanamycin, and then used to inoculate 50-100 mL of liquid LB containing 50 μg/mL kanamycin and 34 μg/mL chloramphenicol. Following overnight incubation at 37oC with shaking (120 rpm), 30 mL of the starter culture was utilized to inoculate 1 L of Auto Induction Medium (AIM) Super Broth Base including Trace elements (AIMSB02, Formedium) containing 2% glycerol and both antibiotics. Bacteria were grown for 72 h at 18 °C in an orbital shaker (150 rpm) and eventually collected by centrifugation at 5000 rpm in a Sorvall H6000A/HBB6 swinging-bucket rotor for 15 min at 4 °C.

Bacterial pellet was re-suspended in 100 mL of the lysis buffer (20 mM Tris-HCl pH 7.5, 200 mM NaCl, 10 mM imidazole, 10 mM 2-mercaptoethanol, 1 mM phenylmethylsulfonyl fluoride (PMSF), 0.02 μM pepstatinA, 0.02 μg/ml chymostatin, 0.006 μM leupeptin, 20 μM benzamidine hydrochloride), incubated with lysozyme (50 μg/mL; Roth) for 30 min in a cold cabinet with head-over-tail rotation, and then broken in the EmulsiFlex-C3 High Pressure homogenizer at 1500 Bar. Homogenate was centrifuged in a Sorvall WX ULTRA SERIES ultracentrifuge (F37L rotor) at 33000 rpm for 45 min at 4°C.

The extract (supernatant after high-speed ultracentrifugation) was used for protein purification using the ÄKTA Xpress system (GE Healthcare), employing nickel affinity chromatography on the 5 mL column compatible with ÄKTA, which was manually filled with Ni-NTA Superflow resin (Qiagen). The column was equilibrated with 25 mL of low-salt (LS) buffer (20 mM Tris-HCl pH 7.4, 200 mM NaCl, 10 mM imidazole, 10 mM 2-mercaptoethanol) prior to extract loading. After protein binding, the resin was sequentially washed with 40 mL of LS buffer, 25 ml of high-salt (HS) buffer (20 mM Tris-HCl pH 7.4, 1 M NaCl, 10 mM imidazole, 10 mM 2-mercaptoethanol), and again with 20 mL of LS buffer. Bound proteins were recovered by elution with 30 mL of buffer E (50 mM Tris-HCl pH 7.4, 200 mM NaCl, 300 mM imidazole, 10 mM 2-mercaptoethanol). Further protein purification was achieved by performing size-exclusion separation of 5 mL of the eluate from the affinity chromatography step on a Hiload 16/60 Superdex S200 column (GE Healthcare), with the use of 1.2 column volumes of gel-filtration (GF) buffer (Tris-HCl pH 7.4; 150 mM NaCl), followed by ion-exchange chromatography on a Resource Q 1 mL column (GE Healthcare), using linear gradient of NaCl (150 mM-1M) in Tris-HCl pH 7.4, 1 mM DTT.

Two fractions corresponding to the maximum of A280nm absorbance were collected after ion-exchange separation, pooled together, mixed with glycerol (30% v/v), aliquoted, snap-frozen in liquid nitrogen and stored at -80 °C. Purified *Tt*Xrn1 was inspected in 10% SDS-PAGE stained with Coomassie Brilliant Blue R-250, along with commercial yeast Xrn1 (NEB; M0338) as a positive control (Supplementary Figure X). PageRuler Prestained Protein Ladder, 10 to 180 kDa (ThermoScientific), was used as a molecular weight marker during electrophoresis. Enzyme specificity towards 5’-monophosphorylated termini was tested by analyzing efficiency of degradation of synthetic 30-mer oligoribonucleotide substrates (5’-ACUCACUCACUCACCAAAAAAAAAAAAACC-3’) bearing fluorescein amidite (FAM) at the 3’-end, and either 5’-monophosphate (5’P-RNA30-FAM-3’), 5’-hydroxyl (5’HO-RNA30-FAM-3’) or 5’-Gppp (5’Gppp-RNA30-FAM-3’) in 1x NEBuffer 3 (NEB; B7003), in 20% denaturing polyacrylamide gels, followed by fluorescence scanning in Typhoon™ FLA 9500 biomolecular imager. Furthermore, the ability to eliminate 18S and 25/28S ribosomal RNAs from total yeast/human RNA samples was examined by treating 1 μg of respective RNA with TtXrn1 for 30 and 60 minutes and running degradation products in 1% agarose gel in 1x TBE containing ethidium bromide. In all biochemical analyses, parallel reactions were carried out using equivalent amounts of commercial Xrn1 as positive control.

### Xrn1 *in vitro* digestion of the pA^+^ fraction

To remove uncapped mRNAs the 35 ng of pA^+^ fraction was digested with home-made *Thermothelomyces (Myceliophthora) thermophilus* Xrn1 *in vitro* at 37 °C in Neb3 buffer (B7003S) for 1 hour in a total volume of 20 μl. The sample was then inactivated 10 min at 80 °C and the RNA was extracted as described above and precipitated.

### Reverse transcription and quantitative PCR analysis

cDNA synthesis was done using SuperScript IV reverse transcriptase form LifeTechnologies (18090050). The reaction mix contained random hexamers (final concentration 2.5 ng/μ), oligo-dT_(18)_ (final concentration 5 μM) as primers and 0.5 mM dNTPs. 1 ug of total RNA or up to 100 ng of pA^+^ fraction was used as template and denatured together with oligo-dT and random hexamers 5 min at 65 °C. After cooling the sample on ice for 2 min, the reaction mix was complemented with buffer, enzyme and RiboLock RNase inhibitor (LifeTechnology; EO 0382) and incubated for 20 min at room temperature and 40 min at 50 °C. The reaction was stopped by incubating samples for 10 min at 80 °C. cDNA was diluted to 400 μl and used. 2 μl of cDNA was used in a total of 5 μl reaction mix for qPCR analysis with commercial SYBR reaction mix [Platinum SYBR™ Green qPCR SuperMix-UDG from LifeTechnology (11733046) or RT PCR Mix SYBR from A&A BIOTECHNOLOGY (2008-1000)] on a LightCycler LC480 Roche apparatus. Data were extracted using the 2^nd^ derivative max method. Oligonucleotide sequences are indicated in Supplemental Table 3.

### RNaseH digestion of *RPS5* 3’end

20 μg of total RNA was mixed with 2 μM RNaseH-targeting oligonucleotide (5’-GGCCAAAGTTTCAGCAATGGTC-3’) in annealing buffer (50 mM Tris-HCl, 50 mM KCl, Ph 8.3) in a total volume of 12 μl and incubated for 2 min at 85°C to be then slowly cooled to 37°C. Where indicated, 2 μM of oligo-dT_(18)_ oligonucleotide (5’-TTTTTTTTTTTTTTTTTT-3’) was also included in order to target poly(A) tail trimming by RNaseH. Next, 8 μl of a mix preheated to 37°C and containing 2.5 U of RNase H (New England Biolabs), 2.5× RNase H reaction buffer (1× buffer: 50 mM Tris-HCl, 75 mM KCl, 3 mM MgCl2, 10 mM DTT at pH 8.3), 25 mM DTT, and 4 U RiboLock RNase inhibitor (Thermo Scientific) was added and incubated for 30 min at 37°C. Subsequently, RNA was precipitated by the addition of 100 μl absolute ethanol and 20 μl solution containing 600 mM sodium acetate (pH 5.3), 10 mM EDTA, 5 μg tRNA, and 5 μg glycogen and incubated at −20°C. Sample pellets were washed with 70 % ethanol and resuspended in RNA loading buffer (formamide with 10mM Tris-HCl pH 8.0, 5 mM EDTA, 0.02 % xylene cyanol).

### Northern blotting

RNA samples were separated on 6% urea-polyacrylamide gels by electrophoresis, transferred to Hybond-N+ membrane (GE Healthcare), and hybridized over-night at 50 °C in ULTRA-Hyb Oligo Hybridization buffer (Invitrogen AM8663) with 1 pmol of 5’ terminally 32P-labelled DNA oligonucleotide probe for RPS5 (5’-CTTAACGGTTAGACTTGGCAACACGTTCCAATTCATCCTTCTTCTTGATAGCGTAAGAAG-3’). The membrane was washed four times with 2XSSC buffer (300 mM NaCl, 30 mM trisodium citrate, pH 7.0) containing 0.5% SDS, each time rotating for 30 min at 42 °C, and exposed to phosphorimager screen. The upper part of the membrane was subsequently cut out, hybridized with a labeled DNA probe for SCR1 (5’-GGTCACCTTTGCTGACGCTGG-3’), and washed and exposed as above. Phosphorimager scans were processed and quantitated with ImageJ software. The indicated pA tail lengths were approximated from DNA size markers.

### Bulk poly(A) tail length analysis

500 ng of oligo(dT)-selected poly(A)+ RNA was subjected to 3’-end labeling in 20 μl of reaction mixture containing: 2 μl of 10x Reaction Buffer for T4 RNA Ligase (Thermo Scientific), 2 μl of 10 mM ATP, 2 μl of 100 mM DTT (Invitrogen), 2 μl of 100% DMSO (Thermo Scientific), 0.5 μl of RiboLock RNase Inhibitor (40 U/μl; Thermo Scientific), 2 μl of T4 RNA Ligase 1 (Thermo Scientific) and 1.5 μl of [5’-32P]pCp (3000Ci/mmol, 10mCi/ml; Hartmann Analytic), for 1 hour at 37 °C. Enzyme was heat-inactivated at 95 °C for 5 min. Labeled RNA was digested with 0.5 μg of RNase A and 10 U of RNase T1 (both from Thermo Scientific) for 20 min. at 37 °C. Reaction was stopped by addition of an equal amount of formamide loading dye (90% formamide in 1xTBE, 0.03% xylene cyanol, 0.03% bromophenol blue) and incubation at 95 °C for 5 min, followed by flash freezing in liquid nitrogen. 5 μl of the sample was run in denaturing 12% sequencing polyacrylamide/8 M urea/1xTBE gel. 3’-end labeled 20 nt-, 31 nt- and 51 nt-long synthetic RNA oligonucleotides (CCCCACCACCAUCACUUA_3_, CCCCACCACCAUCACUUA_14_, CCCCACCACCAUCACUUA_34_; all from Future Synthesis) were run in parallel to the labeled poly(A) tails as molecular size markers. Following electrophoresis, the gel was transferred onto Whatman 3MM filter paper, dried at 80 °C for 2 hours under vacuum, and exposed overnight to a PhosphorImager screen (FujiFilm), which was scanned using a FLA 9000 scanner.

### Nanopore sequencing

DRS libraries were prepared using Direct RNA Sequencing kit (ONT – Oxford Nanopore Technology, SQK-RNA002) according to the manufacturer’s instructions, using 50–200 ng oligo-dT_(25)_-enriched mRNA from Saccharomyces cerevisiae yeast mixed with cap-enriched or total RNA from other organisms (human, mouse, *A. thaliana*, or *C. elegans*), as described in Bilska et al. (2020). Raw data were basecalled using Guppy (ONT). Raw sequencing data files (fast5) were deposited at the European Nucleotide Archive (ENA; for a list of accession numbers see Supplemental Table 4). The list of all sequencing runs, with appropriate summaries, is shown in Supplemental Table 4.

### Bioinformatic analysis

#### Poly(A) lengths determination

DRS reads were mapped to the custom *S. cerevisiae* transcriptome described in Tudek et al. (2021), or to *S. pombe* transcriptome (PomBase, CDS + introns + UTRs) using Minimap2 2.17 with options -k 14 -ax map-ont –secondary = no and processed with samtools 1.9 to filter out supplementary alignments and reads mapping to reverse strand (samtools view -b -F 2320). All unmapped reads were filtered out and discarded from analysis. The poly(A) tail lengths for each read were estimated using the Nanopolish 0.13.2 polya function and only length estimates with the QC tag reported as PASS were considered in subsequent analyses. Since the replicates strongly correlated with each other, unless indicated differently, samples from the same condition were analyzed together. Analyses of changes in median poly(A) tail length and mRNA abundance were performed using R. Tables with the number of counts, mean, median, geometric mean poly(A) tail lengths and quantiles are deposited at Mendeley data (doi: 10.17632/2j3hh37zzs.1).

#### Binning transcripts by pA-tail length

For each set of samples, transcripts were divided into non-overlapping bins based on their median poly(A) tail lengths (binwidth=5). Each bin contained transcripts that were represented by more than 30 reads. Then for each condition and poly(A) length bin number of transcripts in each bin were plotted (only transcripts that were detected in each dataset was included).

#### Binning reads by pA-tail length

For each set of samples reads were divided into bins of either multiple of 5 or multiple of 30 based on poly(A) tails length. Then for each condition and poly(A) length bin read abundances (normalized to sequencing depth) were plotted.

#### Gene ontology analysis

Gene ontology analysis was performed using the BioMart R package. Only unique transcript markers were selected to avoid overlapping gene groups.

### Modeling of decay and deadenylation rates

#### Normalization of RNA abundance

To produce a decay rate model the sequencing data was first normalized to reduce the impact of unequal library size. *ENO2* transcript counts were removed from the raw datasets as in some datasets they served as an external spike-in for other libraries and the abundance of this transcript often introduced unwanted bias to RNA level quantification. For decay modeling the transcript levels were normalized across chase datasets. First, absent raw counts (NA values) were replaced with 0.01. Second, such corrected raw count numbers were divided by a coefficient derived from the sum of *S. pombe* counts in each library and its relative level to other such sums in each chase dataset. Third, count levels were normalized by library size.

Prior to building a model the degradation and deadenylation patterns were visually inspected as shown in Supplemental Figure 1 and Supplemental Figure 2A-J. This led to the conclusion that, depending on the transcript, decay and deadenylation can follow both a linear and an exponential pattern. When building each model, all values, which were greater than the control were removed. This is justified by the fact that regardless of the condition transcription is down-regulated, but not fully switched-off, therefore an increase in mRNA abundance or pA-tail length could indicate *de novo* transcription. Visual inspection of the data also allowed us to determine the time points at which the dataset was reliable enough to model each factor. In the later time points of export block the decreased read number for most mRNAs resulted in an overrepresentation of hyperadenylated transcripts, which is due to the nuclear phenotypes related to depletion of Mex67-AA. Similarly, past the 15 min time point of heat-chase most transcriptionally down-regulated mRNAs were starting to display bi-modal pA-tail distribution, which was indicative of initiation of transcription at loci coding for those mRNAs.

#### Exponential modeling

The exponential model follows the y = e^-kt^ formula, where e = Euler’s constant, t = time and y = fraction of control RNA abundance or pA-tail length. The decay constant k was estimated from the experimental data. Decay and deadenylation constants calculated independently for each dataset were averaged between datasets.

#### Linear modeling

Linear model follows the y = ax + b, where a – decay or deadenylation rate (direction of the slope), b = y value at which x = 0 min (intercept), x = time, y = fraction of control RNA abundance or pA-tail length. Linear modeling was performed using two approaches.

The first method used was linear regression, which for best accuracy was produced on all three replicates combined. In this case, additional data filtering was performed in which all data points displaying an increase in values higher than 1.5-fold at time points equal or larger than 20 min in relation to the preceding time points in the given replicate were removed. Decay and deadenylation rates were derived from a weighted linear regression. Weights were added to the control samples to match the number of values used for modeling decay and deadenylation. This was necessary since the time points close to 0 min were underrepresented causing the best fit curve to greatly diverge from the control sample. Only decay and deadenylation rate estimates were taken for further comparison for which the intercept was close to 1.

Since linear regression produced fewer reasonable best fit curves compared to the exponential fit, we modified the approach to produce an alternative linear model. In the revised approach the lines were fitted to two points: control and time x and the slope factor was averaged within each replicate and then next between replicates. As follows in all cases the intercept was equal to 1.

For both the exponential and linear models if the decay factor could not be determined within the dataset and the mRNA was within the most highly expressed mRNA group [log2(mRNA abundance)>(−10)] it was assigned a random half-life between 60 and 120 min. In such case this is indicated.

The decay and deadenylation rates together with supporting parameters for each model are listed in Supplemental Table 1, corrected values are specified in the table together with the biotype and susceptibility to transcriptional down-regulation during heat stress as defined experimentally by Vinayachandran et al. (2018).

**Supplemental Table S2.** Yeast strains used in this study.

| Name | Genotype | Source |
| --- | --- | --- |
| Wild-type W303 | W303: MAT A; leu2-3,112; trp1-1, can1-100; ura3-1, ade2-1; leu2-3,112; his3-11,15 | T. H. Jensen collection Y159; A. Tudek collection Y13 |
| Dcp2-AID-FLAG | As W303, OsTIR1:URA3, DCP2::mAID-6flag::HygR | This study (T. H. Jensen collection Y3798; A. Tudek collection Y45) |
| Dcp2-AID-FLAG Mex67-AA | As W303; tor1-1 fpr1::LEU RPL13-2xFKBP12::loxP-TRP1-loxP OsTIR::URA | This study (T. H. Jensen collection Y3927; A. Tudek collection Y62) |
|  | MEX67:FRB::kanMX DCP2-mAID-6FLAG::HygR |  |
| Pab1-AID-6HA<br>Mex67-AA | As W303, tor1-1 fpr1::LEU RPL13-2xFKBP12::loxP-TRP1-loxP OsTIR::URA MEX67-FRB::KANmx PAB1-mAID-6HA::HygR | This study (T. H. Jensen collection Y3879) |
| *dhh1Δ* | As W303, DHH1::URA3ca | This study (A. Tudek collection Y68) |
| *lsm1Δ* | As W303, DHH1::URA3ca | This study (A. Tudek collection Y69) |
| Mex67-AA | As W303, tor1-1 fpr1::NAT RPL13-2xFKBP12::TRP1 MEX67-FRB::kanMX6 | Euroscarf from Haruki et al., 2018 (T. H. Jensen collection 2618; A. Tudek collection Y39) |
| Wild-type to match<br>Tucker et al., 2001<br>strains | MATa leu2-3,112 trp1-1 ura3-52 his4-539<br>cup1::LEU2/PGK1pG/MFA2pG | Tucker et al., 2001 (R. Parker collection yRP840; A. Tudek collection Y79) |
| *pan2Δ* | MATa leu2-3,112 trp1-1 ura3-52 his4-539<br>cup1::LEU2/PGK1pG/MFA2pG<br>pan2Δ::URA3 | Tucker et al., 2001 (R. Parker collection yRP1619; A. Tudek collection Y80) |
| *ccr4Δ* | MATa leu2-3,112 trp -1 ura3-52 his4-539<br>cup1::LEU2/PGK1pG/MFA2pG<br>ccr4Δ::NEO | Tucker et al., 2001 (R. Parekr collection yRP1616; A. Tudek collection Y81) |
| *pan2Δ ccr4Δ* | MATa leu2-3,112 trp -1 ura3-52 his4-539<br>cup1::LEU2/PGK1pG/MFA2pG<br>ccr4Δ::NEO pan2Δ::URA3 | Tucker et al., 2001 (R. Parker collection yRP1620; A. Tudek collection Y82) |
| Xrn1-AID | As W303, His+ tor1-1 fpr1::LEU RPL13-2xFKBP12::loxP-TRP1-loxP HIS+ OsTIR::URA3<br>XRN1::AID::KANmx | Tudek et al. 2018 (T. H. Jensen collection Y3688; A. Tudek collection Y43) |

**Supplemental Table S3.**
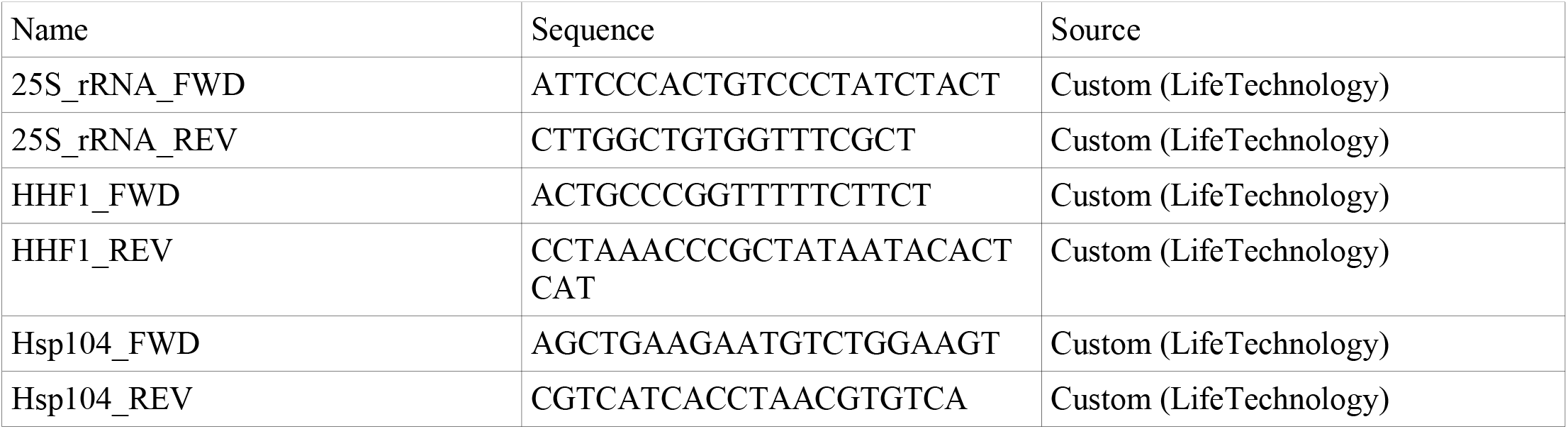
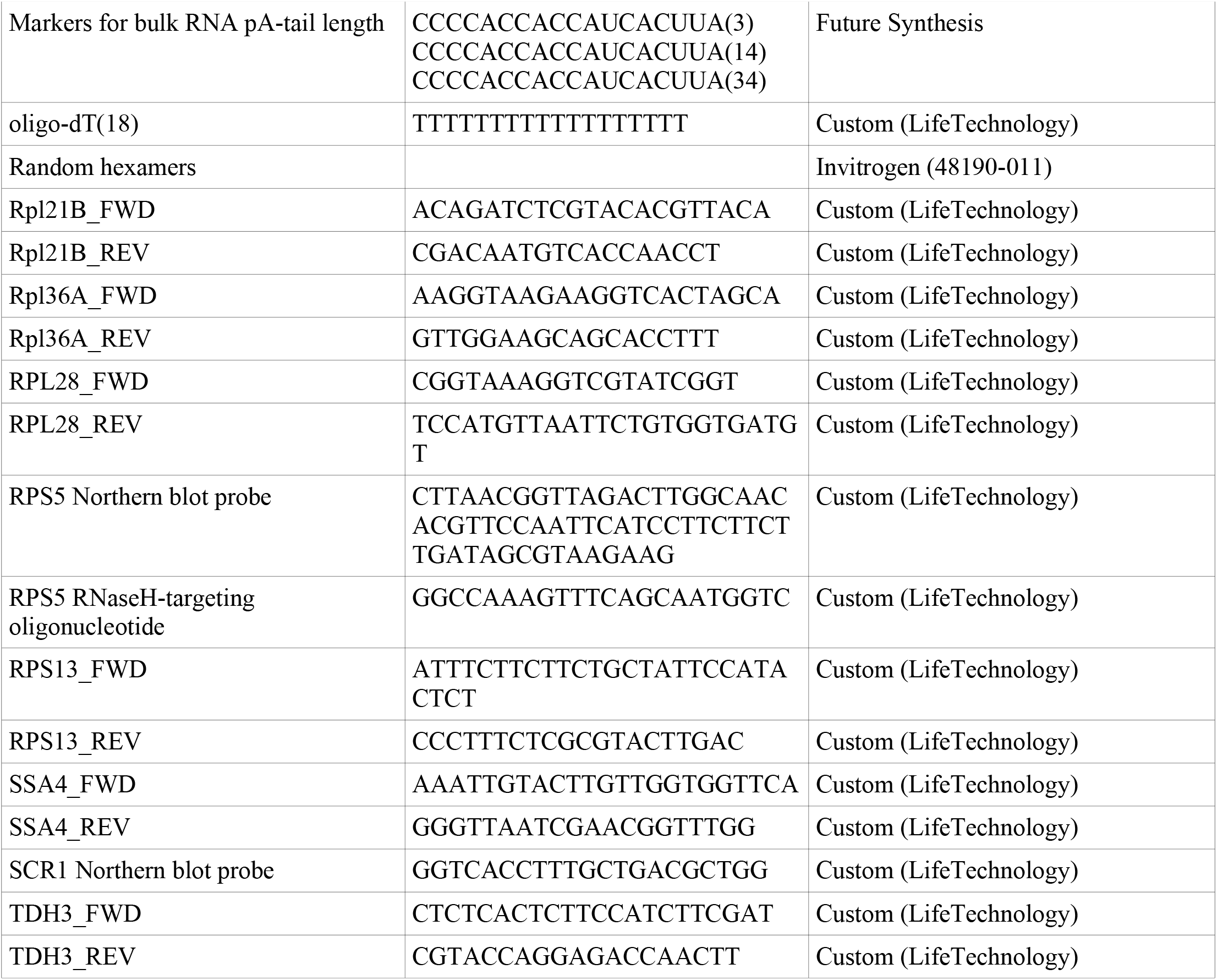
Oligonucleotides used in this study

## Notes

### Competing Interest Statement

The authors have declared no competing interest.

